# Multiple factors regulate i-motif and G-quadruplex structures *in vitro*: analysis of repeated and non-repeated polyG/polyC clusters by circular dichroism

**DOI:** 10.1101/2024.08.26.609788

**Authors:** Levi Diggins, Daniel Ross, Sundeep Bhanot, Rebecca Corallo, Rachel Daley, Krishna Patel, Olivia Lewis, Shane Donahue, Jacob Thaddeus, Lauren Hiers, Christopher Syed, David Eagerton, Bidyut K. Mohanty

## Abstract

The B-form of DNA in the genome contains thousands of sequences that can form various noncanonical structures. Of particular interest are two structures namely G-quadruplex (G4), formed by two or more stacks of four guanine residues in a plane, and intercalating-motif (i-motif, iM) formed by alternately arranged C-C^+^ pairs. Circular dichroism (CD) spectroscopy is a fast biophysical technique to analyze G4s and iMs. We conducted a CD analysis of two types of DNA sequences, one containing tandem repeats and one without, for the generation of G4s and iMs under various environmental conditions, which include pH, buffer composition, boiling, with flanking sequences, complimentary DNA strands, and single-stranded DNA binding protein (SSB). Changes in pH and boiling caused drastic variations in the CD spectra of DNA containing tandem repeats of GGGGCC and GGCCCC from the C9ORF72 gene, although some changes in G4/iM-forming DNA from promoter-proximal regions of several oncogenes also occur. An increase in the number of hexanucleotide repeats generated complex CD patterns at specific pH due to the presence of both G and C bases. The presence of flanking sequences affects CD pattern of a mixture of G4- and iM-forming sequences of the c-MYC promoter-proximal region. SSB disassembled G4 and iMs of all sequences suggesting an *in vivo* role for SSBs in disassembly of G4s and iMs during various DNA transactions.

## Introduction

The genome primarily consists of B-form double-stranded DNA (dsDNA). However, it is interspersed with numerous sequences that can form noncanonical structures including: DNA hairpins, cruciform DNA, triple helices, G-quadruplexes (G4), and intercalating motifs (i-motifs, iMs)^1,2^. Specifically, G4s and iMs are formed by polyguanine (polyG)-rich and polycytosine (polyC)-rich sequences. G4s contain at least two or more stacks of guanine quartets in a planar arrangement joined by Hoogsteen base pairings, whereas iMs contain multiple C-C^+^ base pairs in alternating planes^3,4^. The human genome contains ∼700,000 G4s^5^ and ∼650,000 iMs^6^, many of which are present at key genomic locations, such as telomeres, centromeres, the promoter-proximal regions of numerous oncogenes, and at loci associated with neurodegenerative diseases. G4s and iMs can modulate various transactions on DNA including replication, repair, and transcription; and thus are implicated in cancer and various neurological disorders including amyotrophic lateral sclerosis (ALS), frontotemporal dementia (FTD), Huntington’s disease and others^7–11^.

Because of their role in crucial DNA transactions, G4s, iMs and the factors that regulate their formation and existence are of significance. The factors that regulate G4s and iMs include local pH, ions, temperature, DNA sequences complementary to or flanking G4 or iMs, and various DNA binding proteins such as helicases. G4- and iM-forming sequences are present at: telomeres^12,13^, the promoter-proximal regions of several oncogenes including BCL-2^14^, c-MYC^15,16^, HIF 1-α^17^, EGFR^18^, VEGF^19^, h-RAS^20,21^, and in specific genes associated with various neurological disorders, and are being extensively studied for their generation and biological roles^22,23^. One of the most widely used techniques to study G4s and iMs is circular dichroism (CD) spectroscopy^24^. However, all of these sequences have been analyzed in isolation and in some cases under limited biochemical conditions.

In this study we have conducted a comprehensive and comparative analysis of several G4- and iM-forming sequences under various *in vitro* conditions including pH, buffer composition, salt concentration, effects of complementary DNA sequences, and other factors including a single-strand DNA binding protein (SSB). We have analyzed several G4- and iM-forming sequences from promoter-proximal regions of BCL-2, cMYC, EGFR, HIF-1α, hRAS, and VEGF. BCL-2 (B-cell CLL/lymphoma 2) transcription^14^ is regulated by P1 and P2 promoters, of which the P1 promoter containing G4- and iM-forming sequences is responsible for the majority of BCL-2 transcription^25^. MYC oncogene, which is upregulated in most cancers, contains a 27 basepair sequence representing the nuclease hypersensitive element III (NHE III) region upstream of the P1 promoter; present in this sequence are G4- and an iM-forming elements^15,16^. The Epidermal Growth Factor Receptor (EGFR) gene codes for a transmembrane tyrosine kinase receptor involved in several signaling pathways that promote carcinoma survival and proliferation and is overexpressed in lung adenocarcinomas, breast cancers, and glioblastoma multiforme, making it a target for various cancer therapies^18^. Hypoxia-inducible factor 1 (HIF-1) is a transcription factor involved in the signaling during hypoxia via the expression of dozens of genes^17^. The hRAS oncogene is a GTPase in the RAS family heavily involved in regulating cell division and proliferation and the gene is mutated in up to 30% of human carcinomas, including lung, colorectal, and pancreatic cancer and are typically associated with a poorer prognosis^21^. While some anti-cancer therapies have been developed that target RAS^26^, these therapies generally focus on downstream effectors of the pathway^21,26^. Germline mutations in the hRAS are associated with Costello syndrome, which can result in coarse facial features, intellectual delay, cardiomyopathies, difficulties feeding, larger than average birth weight, and nasal papillomas^27^. hRAS has two major transcription start sites (hRAS1 and hRAS2) that are GC rich and can form G4 secondary structures^20^. The Vascular Endothelial Growth Factor (VEGF) and its receptor, (VEGF) are possibly the most important factors in both pathological and physiological angiogenesis^19^.

We have compared the CD patterns of the G4- and iM-forming sequences of the above-mentioned oncogenes with CD patterns of the sequences containing 2, 4, 6 and 14 tandem repeats of GGGGCC and GGCCCC present at intron 1 of the gene C9ORF72. The intron contains ∼20 copies of the hexanucleotide repeats (GGGGCC/GGCCCC) which is expanded to hundreds or thousands of copies in a significant population of ALS and FTD patients. Although appropriate pH was found to be the most important factor to generate G4s and iMs, other factors tested such as complementary DNA sequence and temperature are shown to play significant roles in their maintenance. We also show here that *E. coli* SSB, which is known to bind to linear ssDNA, reduces the CD peaks of both G4s and iMs in all samples assessed without shifting the wavelength of the peak formation. The results suggest that SSB unfolds the secondary structures, and it may do so during various transactions on DNA including replication and repair. Since the hexanucleotide repeats of C9ORF72 gene contain GGGGCC or GGCCCC, the mix of both ‘G’s and ‘C’s in the hexanucleotide repeats affect G4 and iM formation significantly due to G-C base pairing as the length of the sequences increases and as the environmental conditions change.

## RESULTS

### Repeated vs non-repeated sequences

The C9ORF72 intron 1 contains 20-25 repeats of GGGGCC/CCCCGG (**Fig. 1 A**) which is expanded extensively in patients with ALS and FTD pathogenesis^8,28,29^. Whereas GGGGCC repeat in the sense strand can form G4, GGCCCC in the antisense strand can form i-motif (iM). In contrast to the tandem repeats of hexanucleotide sequences of C9ORF72, the promoter-proximal regions of various oncogenes contain polyguanine/polycytosine clusters which do not contain tandem repeats, nor do they contain specific interrupting nucleotides (**Fig. 1 B** and **C**). All these polyG-rich DNA sequences can form G4s and the polyC-rich sequences can form iMs (**Fig. 1 D** and **E**). To determine the effects of buffer composition and pH on G4 and iM formation, we used five different buffers at four different pHs – sodium cacodylate.KCl at pH 5.5, Tris.acetate at pH 6.0, MES at pH 6.5, Tris.HCl/KCl at pH 7.4 and sodium cacodylate.KCl at pH (7.4) (see Materials and Methods).

**Figure 1:**
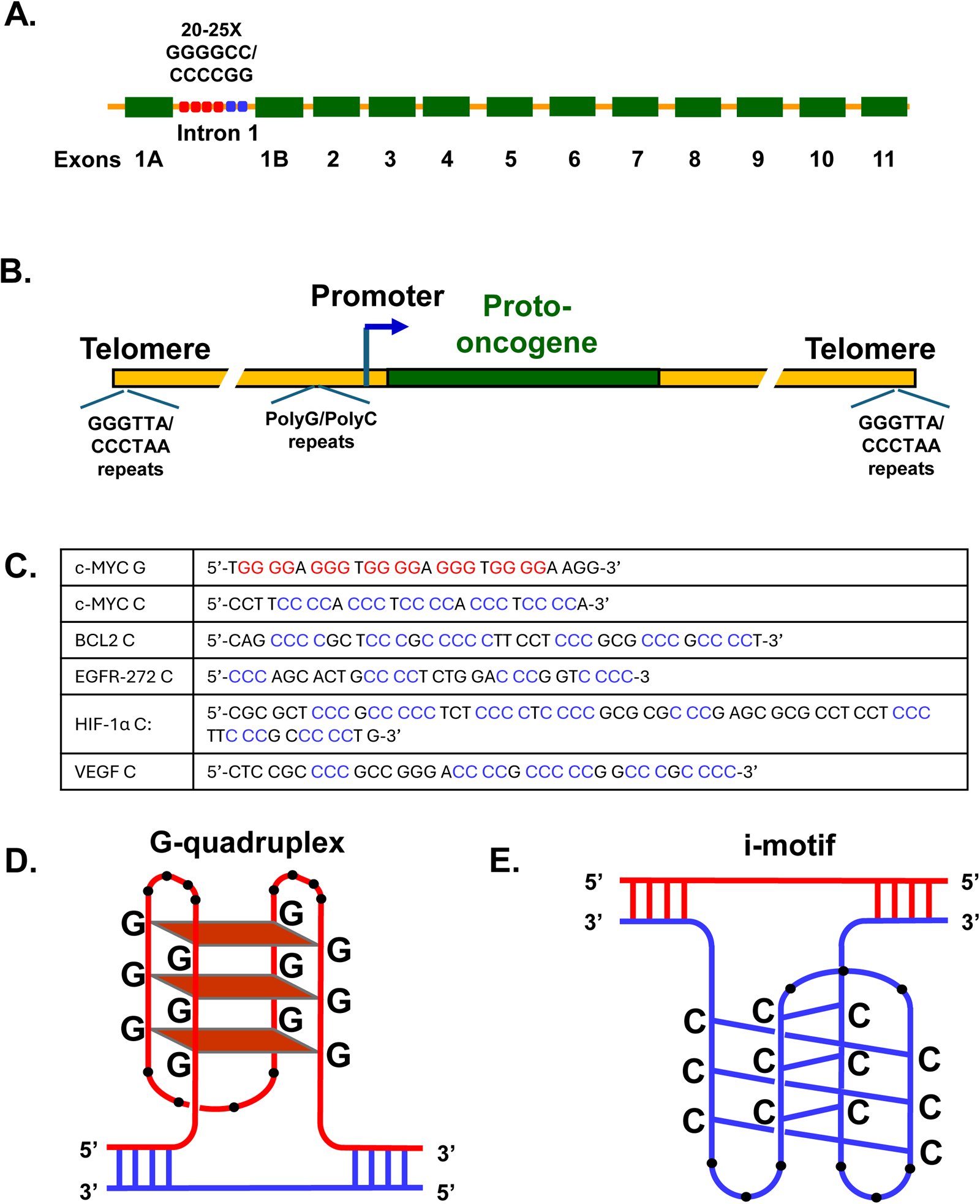
**A.** Hexanucleotide repeats in intron 1 of C9ORF72. **B.** Promotor proximal region. **C.** DNA sequence of promoter-proximal regions of various oncogenes. **D.** Model of G-quadruplex. **E.** Model of i-motif.

### Effects of pH, buffer composition and temperature on iM and G4 formation by the oligodeoxynucleotides containing GGGGCC and CCCCGG repeats

We tested four DNA oligos containing 2, 4, 6 or 14 repeats of 5’-CCCCGG-3’ (C4G2) for their CD patterns in each of the five buffers and either boiled and cooled them or did not boil them at all (unboiled). The CD spectra of all four oligos showed pH dependence (**Fig. 2 C-J**). Typical CD spectra of iM display positive peaks around 288 nm and negative peaks around 262 nm^23,30^. At pH 5.5, the population of iM varied dependently on the sequence length as indicated by the changes in maximum ellipticity. An oligo containing two repeats of 5’-CCCCGG-3’ (C4G2)_2_ showed a strong positive peak around 286 nm, a small positive peak at around 262 nm (where a negative peak should appear), and a negative peak at around 241 nm in sodium.cacodylate pH 5.5 buffer (**Fig. 2 C**). After boiling and cooling (**Fig. 2 D**), there were no major changes in wavelength and the ellipticity showed minimal differences. However, each of the two oligos, (C4G2)_4_ and (C4G2)_6_ whether boiled or non-boiled, showed a positive peak at around 288 nm and a negative peak at around 263 nm (**Fig. 2 E-F and G-H**). Interestingly, boiling of the two oligos followed by cooling resulted in a reduction in iM peak population. We then conducted CD analysis of an oligo containing 14 repeats of the C4G2 hexanucleotide (C4G2)_14_, which is close to the normal ∼20 repeats in the genome. A theoretical analysis of this sequence in iM seeker^31^ predicted it to contain 3 (intrastrand) iMs (**Fig. 2 B**). The oligo (**Fig. 2 I-J**) displayed a positive peak at 284 nm, but it presented a slow and smooth decrease in spectra in its corresponding low pH (5.5 and 6.0) buffers rather than a clear negative peak. In contrast, the higher pH buffers (7.4) showed clear negative peaks at approximately 246 nm. Since each of the hexanucleotide repeats (C4G2)_n_ contains four ‘C’s and two ‘G’s, as the pH increases from 5.5 to 7.4, it is expected that disorganization of iM will occur leading to G-C base-pairing^23^. Accordingly, at pH 7.4, in both sodium cacodylate and Tris.KCl buffers, we consistently observed a shoulder (bump) on the iM spectra at 264 nm where a negative peak is generally observed. In conclusion, oligos containing different repeats of C4G2 showed differences in CD spectra as the size increased and as the pH changed.

**Figure 2:**
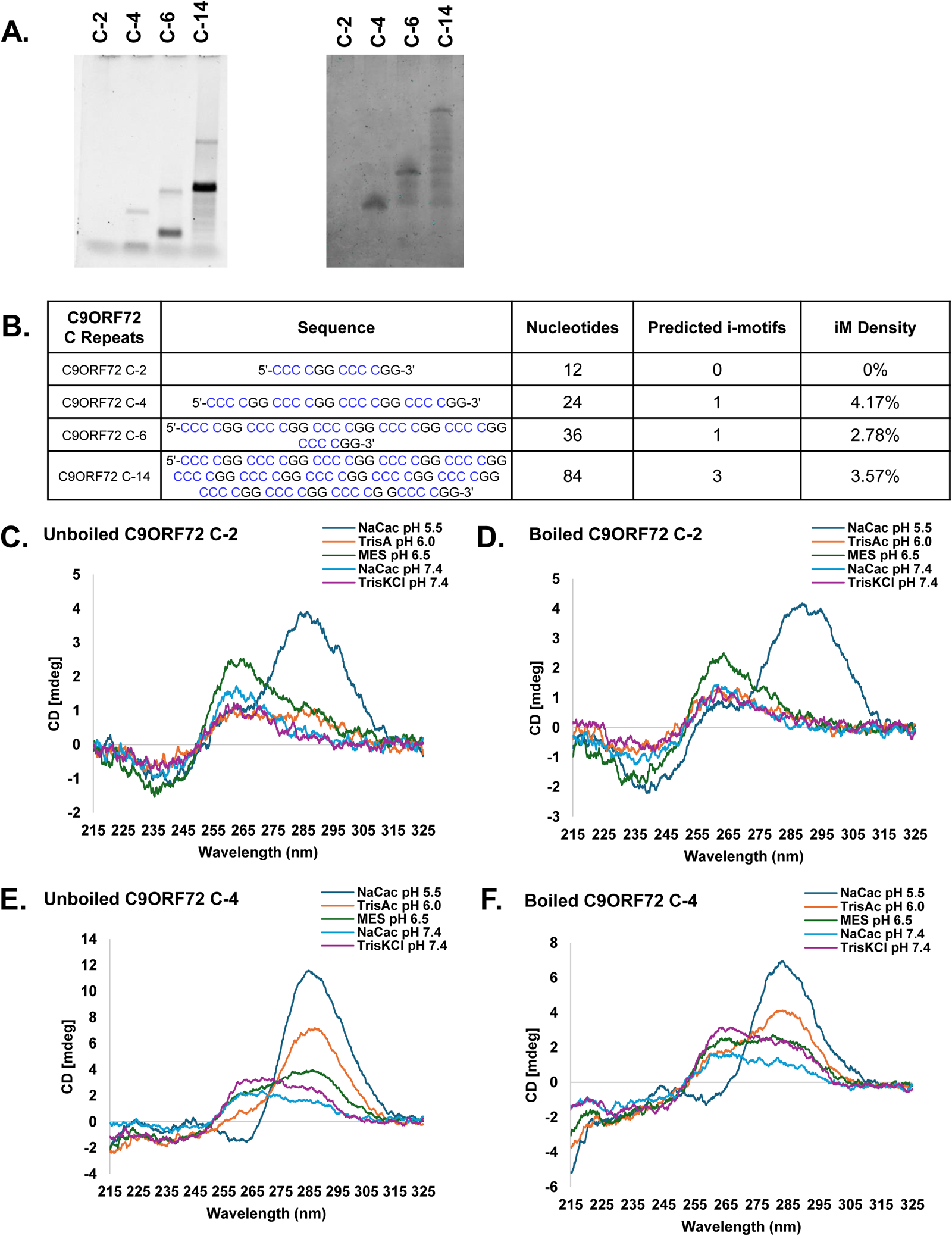
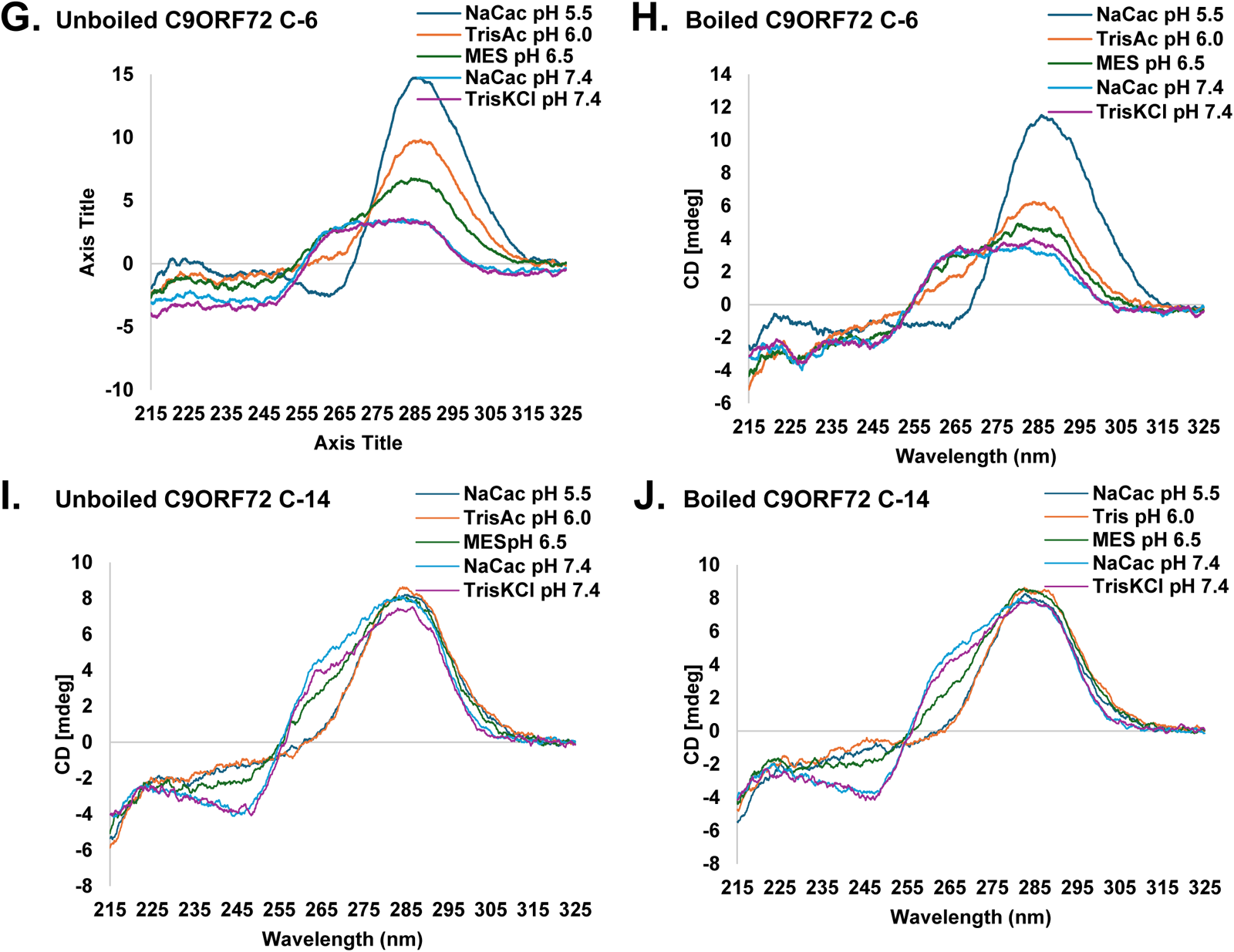
Effects of pH, buffers and temperature on CD spectra of i-M forming sequences of hexanucleotide repeats present at intron 1 of C9ORF72 gene. **A.** 6% polyacrylamide (left) and 15% urea-acrylamide gel (right). **B.** Prediction of iM formation using iM Seeker. **C-J.** CD Spectra

G4 formation is favored at neutral to low alkaline pH. We conducted CD analysis of G4 formation in oligos containing 2, 4, 6 and 14 repeats of 5’-GGGGCC-3’ hexanucleotide repeat (G4C2), complimentary to the four oligos containing C4G2 repeats described in the previous section. Typical features of G4 CD spectra are ∼264 nm max, ∼245 nm min (parallel G4), ∼295 max, ∼260 min (G4) and ∼295 max, ∼260 max, ∼245 min ((or 3+1) hybrid)^32^. The resultant maximum ellipticity changes are influenced by sequence length and pH. As shown (**Fig. 3 C**), the oligo (G4C2)_2_ with 2 repeats of G4C2 shows a typical G4 structure with characteristic max and min peaks of 265 nm and 240 nm. After boiling and cooling (**Fig. 3 D**), C9ORF72 G-2 still shows the expected G4 structure, but with elevated ellipticity. Conversely, the (G4C2)_6_ sample (**Fig. 3 G**) exhibits a higher maximum ellipticity than the (G4C2)_14_ sample (**Fig. 3 I**) in all buffers, including Tris.KCl (pH 7.4, Purple). Additional secondary structures are observed at wavelengths between 280-300 nm, present in both (G4C2)_6_ and (G4C2)_14_ samples with similar ellipticity values. After boiling and cooling (**Fig. 3 H**), the sample containing six repeats had a significant increase in ellipticity and a reduction of secondary structure peaks with the exception of the Tris.acetate buffer. The boiling and cooling of the (G4C2)_14_ sample also showed significant changes in ellipticity. MES pH 6.5 buffer had the largest elliptical change by almost tripling its value, while all other buffers only doubled their ellipticity. Across all samples, G4 structures demonstrate stability in all buffers, particularly when compared to i-motif (iM) formation by polyC-rich DNA samples. The (G4C2)_4_ sample (**Fig. 3 E**) shows little change in ellipticity across the five buffers tested. Interestingly, the (G4C2)_4_ sample shows a dominant positive peak around 295 nm and a second, trivial peak at 247 nm. This is possibly indicative of hairpin formation, with negative peaks appearing at 255 and 233 nm. After boiling, the four repeat sample changed to favoring the expected G4 formation with secondary structures becoming more prominent (**Fig. 3 F**). The (G4C2)_6_ sample (**Fig. 3 G**) in Tris-KCl (pH 7.4, Purple) generates the highest concentration of G4 structures, with minimal change in subsequent buffers. The (G4C2)_14_ sample maintains a hybrid G4 structure formation in all buffers except for sodium cacodylate (pH 5.5, Dark blue). Thus, in our experiments the formation of G4 structure on the polyG-rich strand of C9ORF72 displays ellipticity at a wavelength of ∼265nm with a preference for formation in neutral/low alkaline buffers consistent with the literature. The resultant maximum ellipticity changes are dependent on sequence length and pH. The presence of only two hexanucleotide repeats (G4C2)_2_ results in minimal formation of G4 structure in all buffers with the (G4C2)_6_ sample producing a higher maximum than (G4C2)_14_ in all buffers including Tris.KCl (pH 7.4, Purple). The formation of other secondary structures occurred at a wavelength 295-305 nm that is present in both the (G4C2)_6_ and (G4C2)_14_ samples with a similar ellipticity. Across all samples (G4C2)_4_ displays stability in all buffers, especially in comparison to iM formation in the cytosine strand. In the (G4C2)_4_ sample there is little to no change in ellipticity through all 5 buffers. In conclusion, the (G4C2)_6_ sample in Tris.KCl (pH 7.4, Purple) generated the highest concentration of G4 structure with little to no change in subsequent buffers and G4 in C9ORF72 intron 1 are more stable across a wide range of pH in comparison to their iM-forming counterparts.

**Fig. 3.**
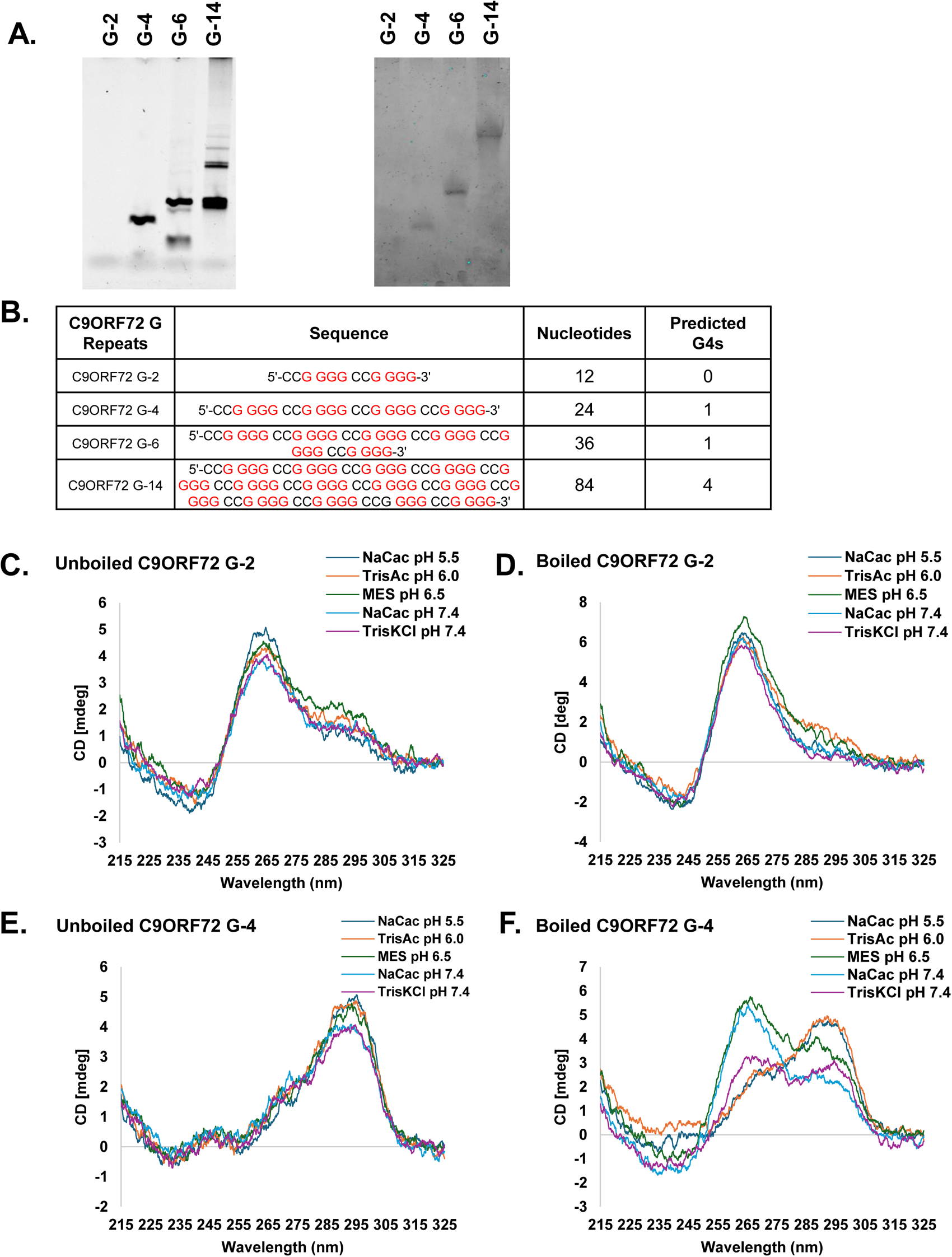
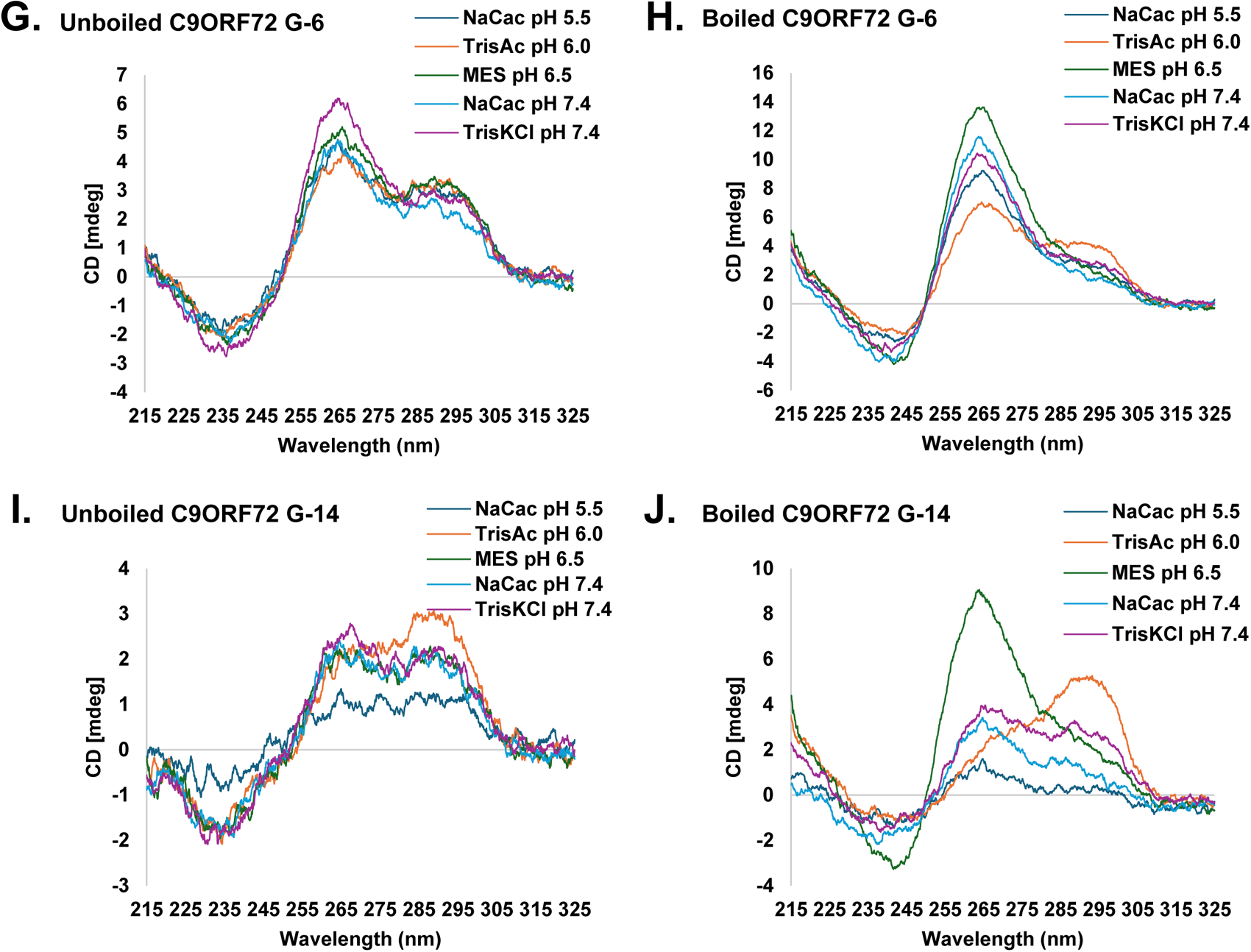
Effects of pH, buffers and temperature on CD spectra of C9ORF72 G-strand oligos (G2, G4, G6 and G14). **A.** 6% polyacrylamide gel (left) and 15% urea gel (right) **B.** Prediction of G-quadruplex formation using QGRS Mapper **C-J.** CD Spectra

### Effects of pH, buffer composition and temperature on iM formation by polyC-rich sequences present at oncogene promoter-proximal regions

Several oligonucleotides – both G-rich and C-rich – representing DNA sequences from oncogene promoter-proximal regions were studied for iM and G4 formation in the five different buffer- and pH-conditions as described above. The CD spectra for the polyC-rich DNA samples (BCL2, c-Myc, EGFR, HIF-1a, hRAS and VEGF) across these different conditions show significant variations in peak ellipticity among them, as well as from C9ORF72 (CCCCGG)_n_ DNA sequences. The unique pH requirements for iM formation in each oncogene highlight their distinct possible regulatory roles. Each DNA sample was entered into iM Seeker to predict the number of i-motif structures based on the sequence (**Figure 4 B**) and compared with our data.

**Fig. 4.**
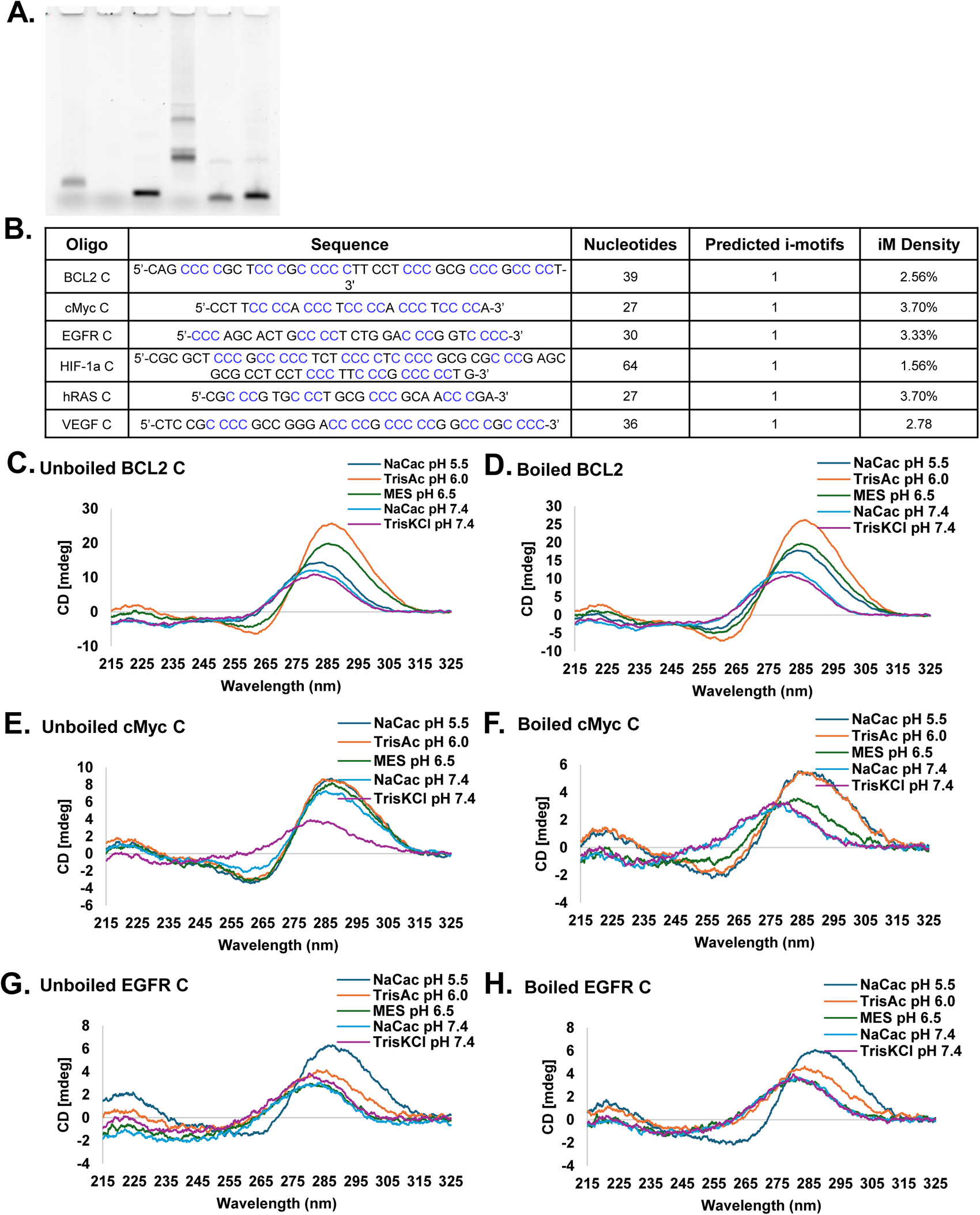
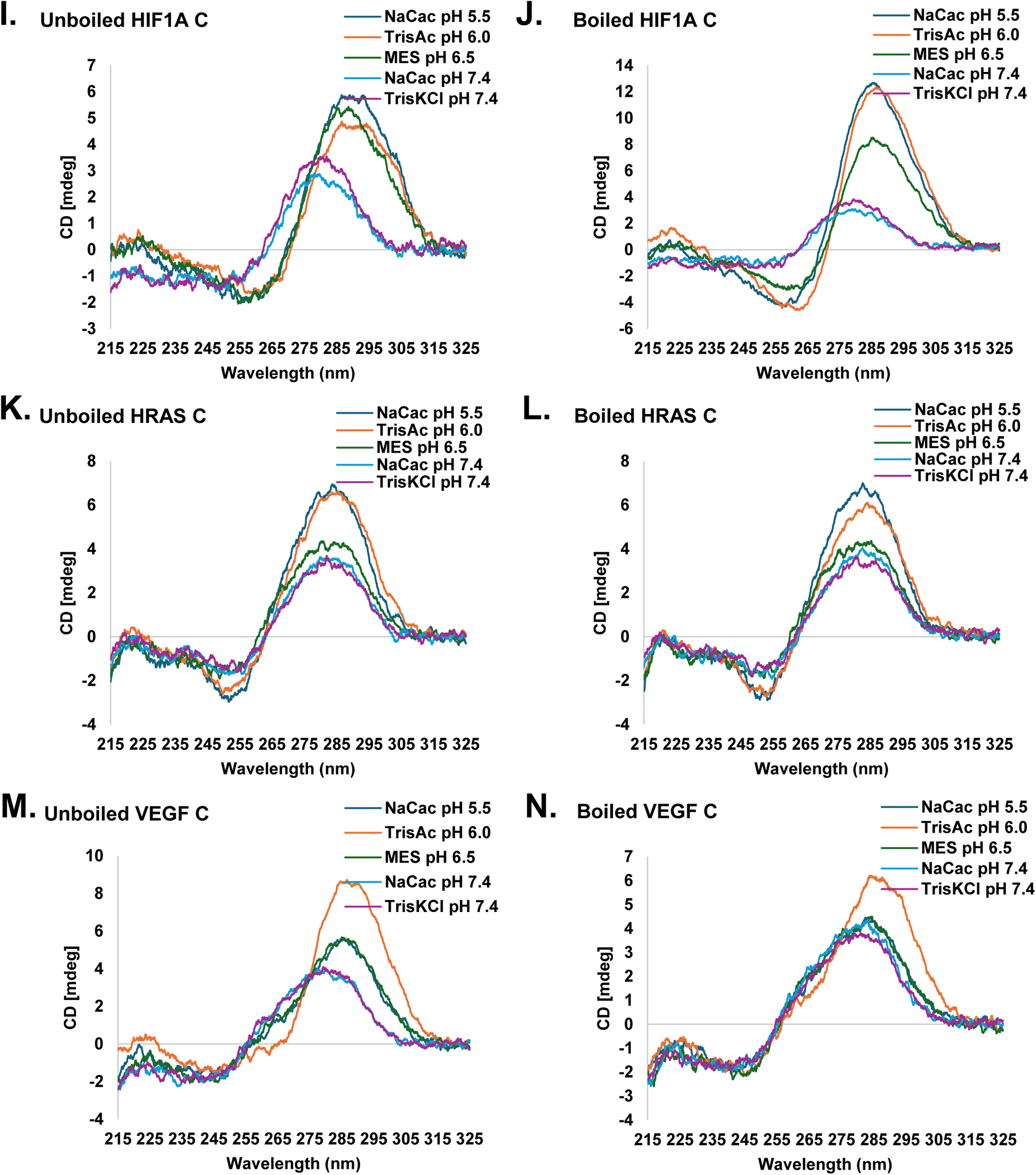
CD spectra of C-strand oligos of oncogene promoter sequences. **A.** 6% polyacrylamide gel **B.** Prediction of i-motif formation using iM Seeker **C-N.** CD Spectra

The BCL-2 oncogene affects apoptosis and contributes to tumorigenesis such as leukemia/lymphoma^14^. A 39-nucleotide long oligo from the BCL-2 promoter-proximal region was studied in all five buffers (**Fig. 4 C**) for its iM properties. The absolute peak ellipticity is the highest in the BCL-2 C sample, compared to all other oncogenes. Formation of the iM structure was highly favored in the Tris.acetate buffer (pH 6.0, Orange) represented by the highest ellipticity, followed by MES buffer (pH 6.5, Green) at ∼285nm. Both Tris.acetate and MES buffers show clear troughs at ∼260nm, aligning with the known iM spectra. All remaining buffers showed a slight peak shift to approximately 280 nm with a lower ellipticity. Na.Cac buffer produces the third highest ellipticity (pH 5.5, dark blue) followed by Na.Cac (pH 7.4, light blue) and Tris.KCl (pH 7.4, purple). After boiling and cooling, the oligo continued the same pattern as the unboiled samples apart from the one in Na.Cac pH 5.5 buffer (**Fig. 4 D**); this buffer showed a shift toward the known iM wavelengths at approximately 284 nm.

The c-MYC proto-oncogene^33^ was analyzed for iM formation under various conditions as for BCL-2 oligo. A 27 base-long oligo representing the nuclease hypersensitive element III (NHE III) region upstream of the P1 promoter of c-Myc was used for CD analysis in five different buffers at all four pH conditions. As expected, and as shown in **Fig. 4E**, the CD spectra of the polyC-rich c-MYC oligo displayed a dominant positive band at 287 nm and relatively weak negative band at 262 nm in Na.Cac buffer at pH 5.5. As the pH increased from 5.5 to 6.0 and 6.5, the peak shifted slightly to a lower ellipticity. In Tris.KCl (pH 7.4, Purple), the positive peak shifts to a lower wavelength of ∼280 nm and becomes less symmetric, suggesting destabilization of the i-motif. All c-MYC C buffer samples showed the traditional iM CD spectra except for Tris.KCl (pH 7.4, purple). The formation of iM is maintained but gradually decreases with the decreasing buffer acidity. The unboiled sample (**Fig. 4F**) formed iM in the same manner at pH 5.5 and 6.0; however, as the pH increased to 6.5, the iM started disassembling and the peak shifted to a lower wavelength. As the pH changed to 7.4, whether in Tris.KCl or Na.Cac buffer, the iM disassembled further and shifted further to a lower wavelength.

The protein encoded by epidermal growth factor receptor (EGFR) gene is involved in cell growth and survival and is generally mutated in lung cancer^18^. The 30 base EGFR oligo sample tends to show lower ellipticity compared to other sequences with a max ellipticity at ∼289 nm at the lowest pH (sodium cacodylate pH 5.5, dark blue) and reduced ellipticity at all other pH conditions. A minor decrease in wavelength can also be observed at all other pH (**Fig. 4 G-H**).

The protein coding gene HIF1A, or hypoxia inducible factor 1-alpha, is a transcription regulator of cellular and system response to hypoxia^17^. The longest oligonucleotide tested, at 64 bases long HIF1-α, showed a similar pattern (**Fig. 4 I-J**) to the c-MYC oligo with some variations. The pattern of the peaks in both sodium.cacodylate and Tris buffers at pH 7.4 were the same. In conclusion, as the pH increased the iM peak decreased and as the pH increased especially from 6.5 to 7.4, the peak shifted from higher wavelength to lower wavelength. The CD pattern of the protein product of HIF-1α gene (**Fig. 4 I-J**) maintains a maximum peak ellipticity in Sodium cacodylate (pH 5.5, dark blue) and exhibits similar peak heights in Tris.acetate (pH 6.5, Orange) and MES (pH 6.0, Green). At a pH of 7.4 there is a significant drop in ellipticity and a decrease in wavelength absorption to ∼280nm. The HIF-1α sample exhibited a maximum ellipticity in MES (pH 6.5, Green) with no significant variations in ellipticity between all buffers.

The hRAS proto-oncogene, which encodes proteins that help regulate cell signaling pathways and cell division, is associated with a wide variety of cancers and other disorders^20,21,27^. As shown (with boiled and cooled or unboiled samples) in **Fig. 4 K-L**, CD analysis of an unboiled 27 nucleotide polyC-rich hRAS sequence produced a maximum peak ellipticity in both sodium cacodylate (pH 5.5, Dark blue) and Tris.acetate (pH 6.0, Orange) buffers at ∼284nm. Ellipticity was reduced at the three remaining pH buffer conditions (as acidity decreased), with no significant changes in wavelength. The troughs of all five samples had a similar wavelength of ∼252nm but the lowest pHs, 5.5 and 6.0, had a lower ellipticity compared to the higher pH samples. After boiling and cooling the hRAS samples, the spectra remained relatively stable compared to the unboiled samples with no major changes in ellipticity or wavelength. The only noticeable change occurred in Tris.acetate buffer, which is a minor reduction of ellipticity. Out of all oncogenes tested, hRAS appears to be the most stable despite the same temperature treatment. VEGF, plays an important role in angiogenesis and has been linked to tumor progression^19^. CD analysis was carried out with a 36 base-long boiled and cooled or unboiled oligo (**Fig. 4 M-N**). The oligo produced a maximum ellipticity in the Tris.acetate buffer (pH 6.0, Orange) at ∼287nm in comparison to all other buffers. There are nearly identical peak heights between both sodium cacodylate (pH 5.5, dark blue) and MES (pH 6.5, Green) buffers at slightly lower wavelength of ∼285nm. These nearly identical ellipticities are also present between sodium cacodylate (pH 7.4, Dark blue) and Tris.KCL (pH 7.4, Purple) buffers but at a further reduced wavelength (∼280nm) and lower ellipticity. While there is no definitive negative peak or trough, all five buffers tend to be at their minimum ellipticity around ∼240nm. Tris.acetate shows a steep decrease in ellipticity from ∼282nm to ∼265nm while all others show a more gradual decrease. After boiling and cooling the samples, the highest peak was still formed in Tris.acetate buffer but expressing a lower ellipticity compared to the unboiled sample. For all other buffers, the wavelength and ellipticity have equalized at ∼282nm with approximately the same ellipticity as the unboiled pH 7.4 samples.

Although there is still no absolute trough, all five buffers have the same negative peak at ∼240nm, but Tris.acetate is more consistent with the other four buffers compared to the unboiled samples.

### Effects of pH, buffer composition and temperature on G4 formation by polyG-rich sequences present at oncogene promoter-proximal regions

The CD spectra of the polyG-rich sequences of BCL-2, c-MYC, EGFR, HIF-1α, hRAS and VEGF across the five different pH conditions show less significant variations in peak ellipticity compared to their polyC-rich counterparts; all samples were either boiled and cooled or not boiled (unboiled) before CD measurement (**Fig. 5**). All sequences were entered into QGRS Mapper to predict the formation of G4s (**Figure 5A**).

**Fig. 5.**
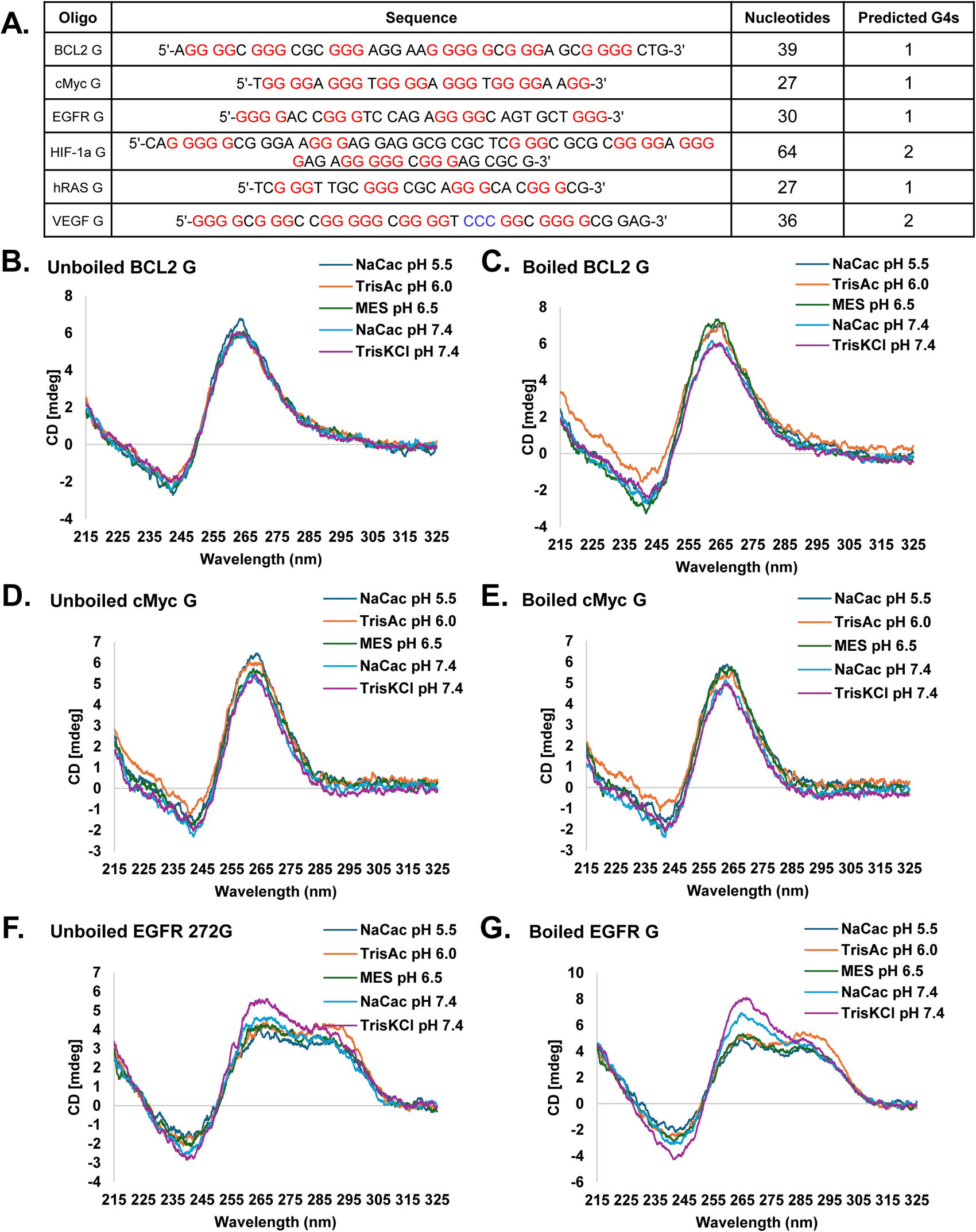
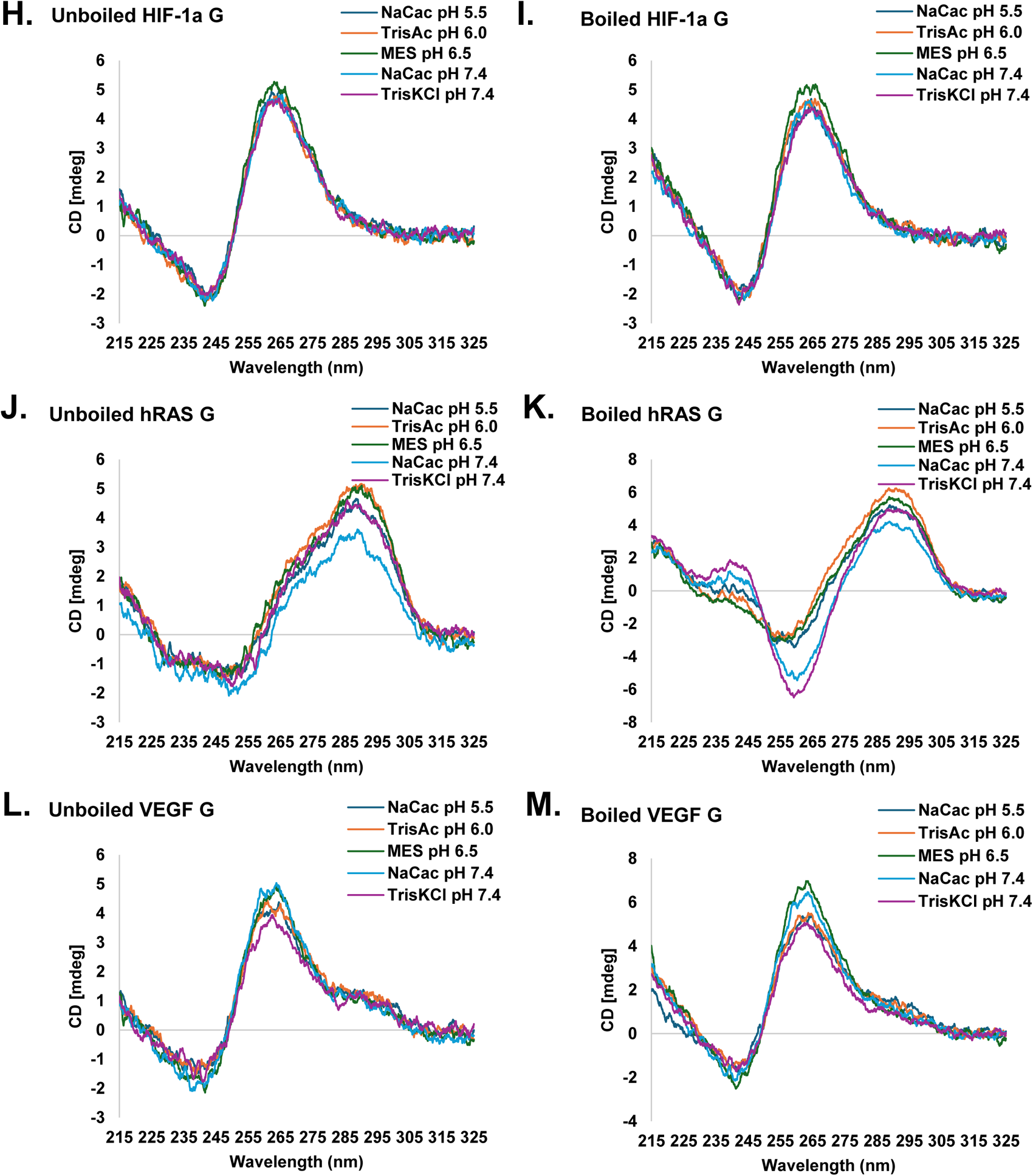
CD spectra of G-strand oligos of oncogene promoter sequences. **A.** Prediction of G-quadruplex formation using QGRS Mapper **B-M.** CD Spectra

The BCL-2 G oligo shows the expected positive and negative peaks for parallel G4s at their respective wavelengths of ∼264nm and ∼244nm (**Fig. 5 B-C**). Sodium cacodylate (pH 5.5) buffer has a slightly elevated ellipticity compared to the buffers at higher pH, but there is no deviation in the pattern from the standard G4 structure. After boiling and cooling the samples, there were minor changes in ellipticity. Parallel structure wavelengths remained unchanged. As with the BCL-2 G oligo, c-MYC G showed very little changes after boiling (**Figure 5 D-E**). Interestingly, the buffer with the lowest pH (sodium cacodylate pH 5.5), displayed the highest peak with ellipticity decreasing with increases in pH. Boiling showed insignificant peak changes. The only minor change was in the case of the c-MYC G oligo in MES pH 6.5 buffer, which slightly elevated ellipticity compared to the unboiled sample. The oligo EGFR G produced one of the most unexpected nonconformant results from the other guanine rich sequences (**Figure 5 F-G**). Interestingly, each of the five buffers produced two positive and one negative peak, with Tris.KCl pH 7.4 generating the highest ellipticity (see Discussion). After boiling, all five pH samples remained the same wavelength structures, but with increased ellipticities. As expected, HIF-1α G produced the conformational peaks associated with G4s (**Figure 5 H-I**). Unlike BCL-2 G and c-MYC G, HIF-1α G displayed the highest ellipticity in both unboiled and boiled samples in MES pH 6.5 buffer. The most unanticipated results were displayed by hRAS G oligo. Contrary to other polyG-rich sequences, the hRAS G oligo produced an iM type peak in all five buffers at 4 pH conditions (**Figure 5 J-K**). Producing a positive peak at approximately ∼290nm and a negative peak around ∼250nm, hRAS G conforms more to iM than G4. There are also vague secondary positive peaks around ∼265nm. Tris.acetate buffer generated the highest ellipticity followed by MES pH 6.5, sodium cacodylate pH 5.5, and Tris.KCl and sodium cacodylate pH 7.4. Boiling the samples created the largest change in negative ellipticity of hRAS G oligo than any other G oligo used. In addition to the ellipticity change, the negative peak also shifted approximately 10nm to ∼260nm. Consistent with the standard G4 CD spectra, VEGF G oligo produced positive peaks at ∼263nm and negative peaks at ∼240nm in all five buffers (**Figure 5 L-M**). The oligo displayed the highest ellipticity in sodium cacodylate (pH 7.4) buffer, while had the lowest ellipticity in Tris.KCl (pH 7.4) buffer. Thus, the buffer composition appears to affect the VEGF G structure despite the same pH. After boiling, ellipticity of VEGF G oligo changed in the sodium cacodylate pH 7.4 and MES pH 6.5 buffers, with MES pH 6.5 showing the highest ellipticity and Tris.KCl pH 7.4 showing the lowest.

In addition to using boiled/cooled and unboiled oligos in all above experiments, we conducted an experiment to measure CD in which temperature was raised from 20 to 65°C. We studied the BCL-2 C oligo for this experiment. The oligo was analyzed in the sodium cacodylate pH 5.5 buffer (**Fig. 6**). As the temperature was raised from 20°C to 65°C, the ellipticity was measured in 5°C intervals. Ellipticity remained constant between 20°C and 35°C before significantly reducing between 35°C and 45°C. Temperatures above 45°C had little effect on ellipticity.

**Figure 6:**
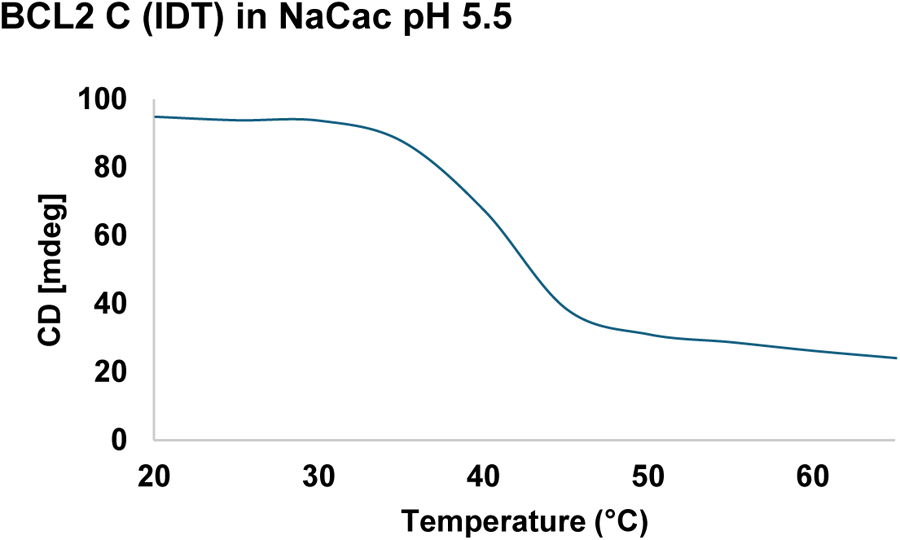
Effect of temperature on iM and G4

### Effects of complementary strands

Despite G4 and iM formation at specific locations genomic DNA contains both strands of DNA everywhere except at telomeres. Therefore, we wished to measure CD pattern of one oligo in the presence of its complementary strand (**Fig. 7**). In two different sets of experiments, c-MYC C oligo was titrated (at three different concentrations) against a fixed concentration of c-Myc G oligo either at pH 5.5 (**Fig. 7 A**) or 7.4 (**Fig. 7 B**). Because of the complementarity between the two oligos, significant interactions occur between the two oligos that should influence the formation of both iM and G4 and this influence is dependent on the concentration of each strand and the cellular pH. At pH 5.5, there was a dose dependent decrease in G4 formation within the c-MYC G sequence with increasing additions of iM forming c-MYC C oligo in an acidic solution. There was a significant change in G4 expression even with a 2:1 ratio (purple) of polyG-rich oligo to polyC-rich c-Myc oligo. This reduction was proportional with each subsequent addition of polyC-rich oligo at both 2:2 (green) and 2:3 (black) ratios. In addition, there was a non-proportional increase in i-Motif formation with each subsequent addition of the polyC-rich oligo. This could indicate a relationship between the formation of iM with the complimentary strand in acidic environments. At pH 7.4, there was a dose dependent increase in G4 formation within the c-Myc G sequence with increasing additions of iM forming c-MYC C sequence in Tris.KCl (pH 7.4) Buffer. The control (light blue) was G4 forming sequence alone which exhibited maximum ellipticity at ∼265nm. The increase in ellipticity was proportional to the concentration of added polyC-rich oligo. A separate iM peak was not present but there was an increase of ellipticity at wavelength >285 nm. This could indicate changes in structural heterogeneity through non-specific interactions or concentration effects. Due to the relatively dynamic nature of the cellular environment, all these data indicate that each structure (G4 or iM) may influence the occurrence of the other and their formation is more likely to occur in specific cellular processes or alterations including disease states that may influence cellular pH (e.g., hypoxia) and DNA replication. It may be noted that the G4 and iM graphs are not displayed independent of each other in these experiments because of the proximity of two graphs. Therefore, our interpretation may be more simplistic than what is really happening in solution.

**Fig 7.**
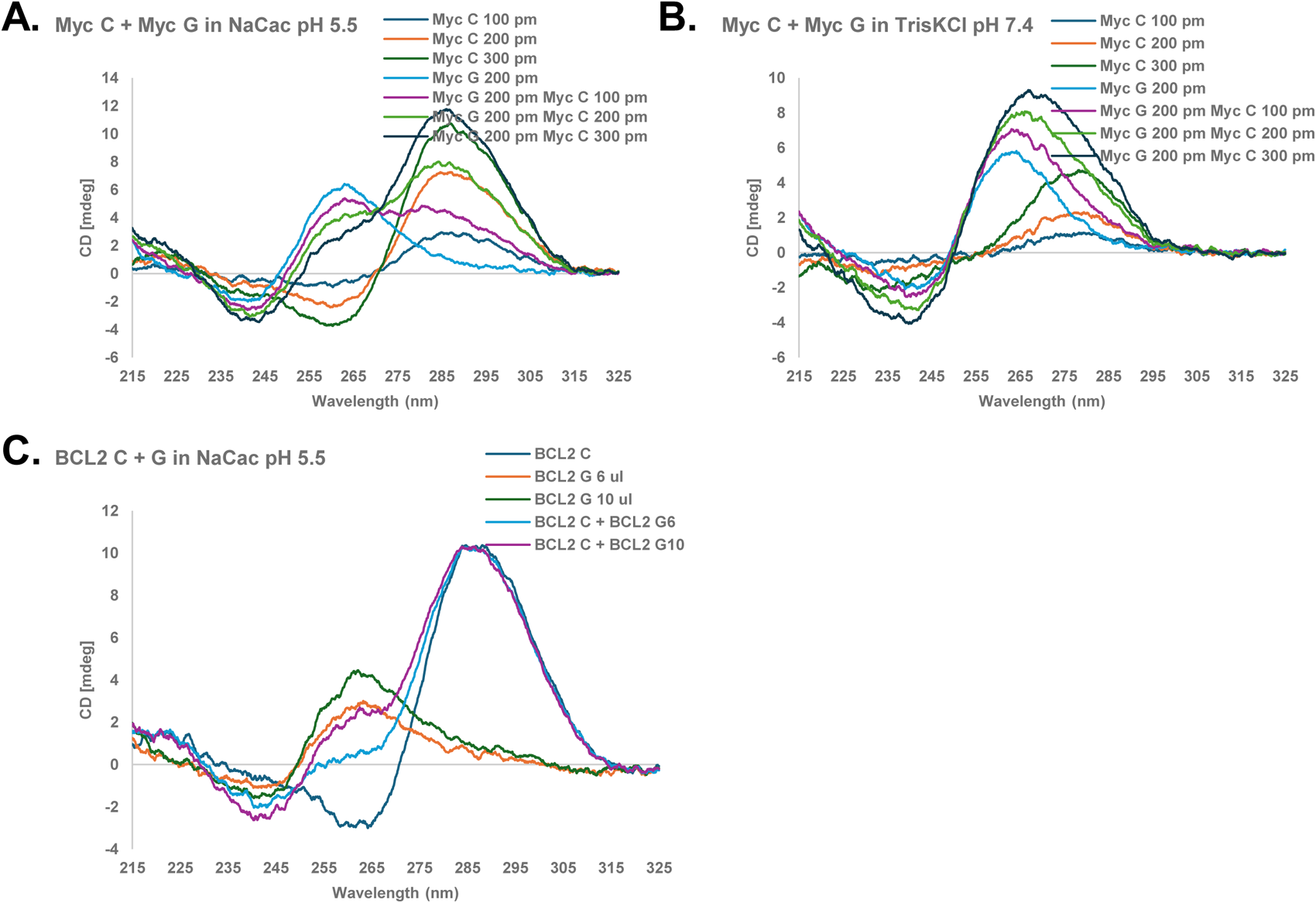
Effects of complimentary strand DNA on G4 and iM A. c-Myc in NaCac pH 5.5 B. c-Myc in TrisKCl ph 7.4 C. BCL2 in NaCac pH 5.5

### Effect of SSB Protein on iM and G4 Structures

DNA replication and repair unwind dsDNA generating ssDNA inside the cell and the ssDNA is protected by coating with SSB in *E. coli* and replication protein A (RPA) in eukaryotes^34–36^. G4s and iMs can be formed during DNA unwinding and various proteins, including helicases, have been demonstrated to unfold the secondary structures^37,38^. We predict that SSB and RPA which bind to ssDNA nonspecifically should help unfold G4s and iMs. To test the hypothesis, we conducted CD analysis of polyC-rich oligos of BCL-2 C, c-MYC C, EGFR, hRAS, and VEGF (**Fig. 8 A-E**). We also used polyG-rich oligos of BCL-2, c-MYC, EGFR and HIF-1α. The formation of G4 and iM structures was influenced negatively by increasing the concentrations of the SSB protein in all samples (**Fig. 8**). BCL-2 C displayed measurable changes in ellipticity with addition of SSB. There are uniform reductions in positive peak height when increased concentrations of SSB were added. Interestingly, there was little change in negative peak height with the additions of SSB. In the c-MYC C sample, there was a reduction of iM with the addition of SSB (Orange) compared to the baseline c-MYC oligo (Dark Blue). There was a concentration-dependent decrease in the iM peak with each subsequent addition of SSB to the sample (Green and Light Blue). Similarly, there was a significant reduction in iM formation by EGFR sequence upon addition of SSB (purple) compared to control (green). In the same fashion, there was a major shift in ellipticity of hRAS C oligo with the addition of SSB. Finally, the formation of iM within the VEGF sequence exhibited a concentration dependent reduction in the presence of SSB.

**Figure 8:**
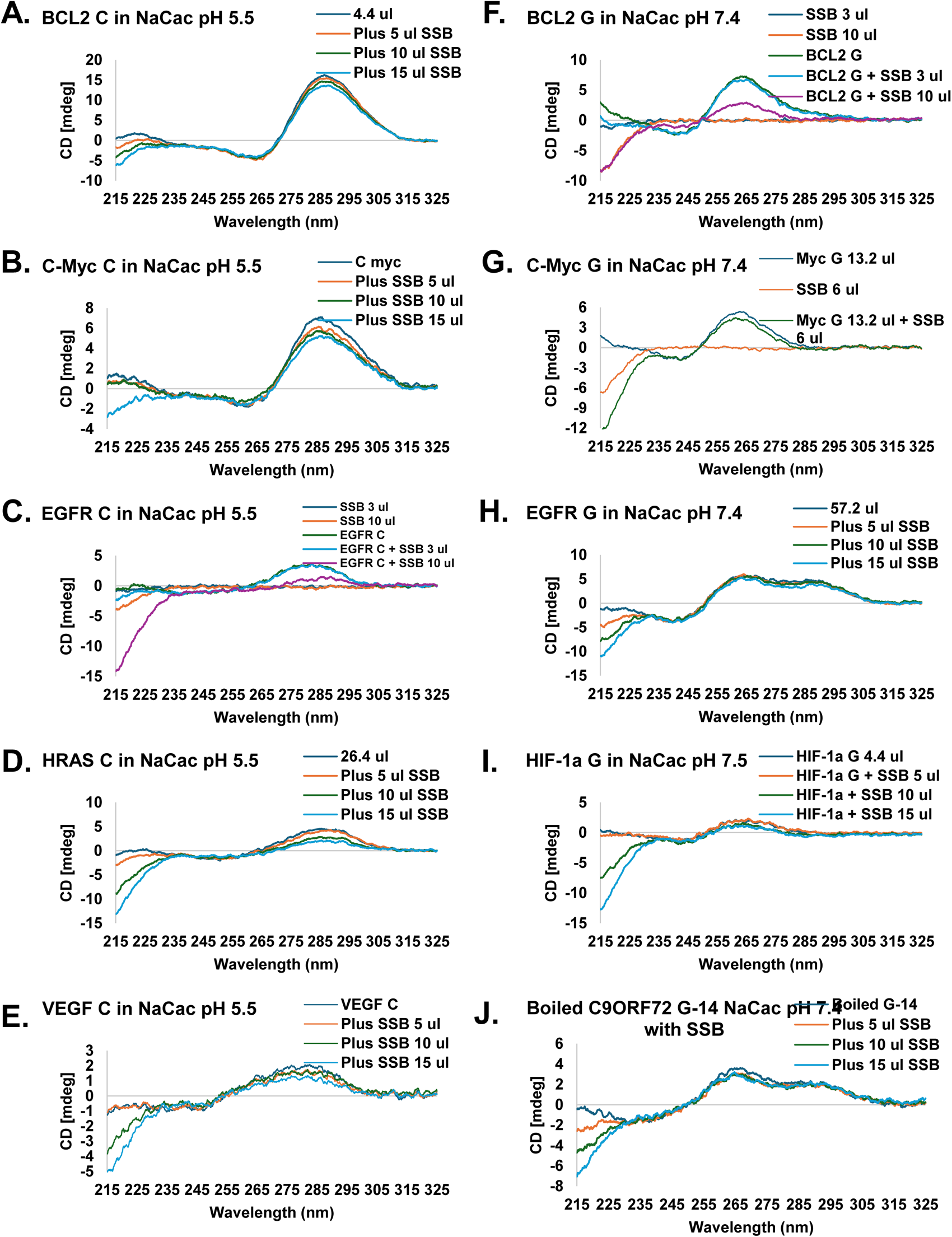
Effect of Proteins

We determined the effect of SSB on G4 formation using four different oligos from promoter-proximal regions of BCL-2, c-MYC, EGFR, and HIF-1α (**Fig. 8 F-I**). The formation of G4 within the BCL-2 G sequence was significantly reduced (purple) by the addition of SSB. Similarly, the formation of G4 within the c-MYC sequence was also reduced with the addition of SSB (green) compared to its control (Dark blue). However, SSB caused a minimal change in the ellipticity of G4 of EGFR G oligo. This is probably due to the large amount of oligo required to achieve the ellipticity shown in the figure and will therefore require a large amount of SSB, which we have not tested. Experiments are underway to test this possibility. Similarly, the G4 peak of HIF-1α G showed minor changes with the addition of SSB.

In conclusion, the i-motif structures exhibited greater sensitivity to SSB, requiring smaller amounts of SSB to reduce iM formation compared to G4. Regardless, the reduction in secondary structure formation by SSB suggests its role in DNA replication and DNA repair.

### Effect of flanking sequences on iM and G4 formation

All G4- and iM-forming sequences are flanked by DNA sequences that generally form linear dsDNA. Therefore, we wanted to determine the effects of flanking sequences on the generation of G4 and iM. For this, we used two complementary oligos from the promoter-proximal region of c-MYC in the middle of which reside the iM- or G4-forming sequences. The flanking sequences on each side contain 20 or 23 nucleotides. **Fig. 9A** shows the extended sequences along with the respective QGRS Mapper and iM Seeker predictions. As shown in **Fig. 9**, the curtailed c-MYC oligos from **Fig. 4E** and **Fig. 5E** were compared to the extended Forward (G) and Reverse (C) strands. In addition to the comparison of the curtailed and extended strands, the extended strands were combined to see the effects of the extended complimentary sequences. **Fig. 9B** shows a significant decrease in ellipticity between the curtailed c-MYC C and flanking Forward (C) sequences, when measured in sodium.cacodylate buffer (pH 5.5). c-MYC C showed a positive peak at ∼285nm and a negative peak at ∼262 nm. The c-MYC C Reverse sample had a positive peak slightly shifted to ∼290 nm and a negative peak at ∼259 nm. In contrast, the curtailed c-MYC G and flanking Forward (G) sequences show minimal ellipticity change. Both have positive peaks at ∼263nm with the flanking Forward (G) sequence having a slightly higher ellipticity. For negative peaks, both have the same wavelength of ∼242 nm but with the flanking Forward (G) sequence having a significantly lower ellipticity. When the Forward (G) and Reverse (C) sequences were combined, two positive peaks formed at ∼283 nm and ∼264 nm, along with a negative peak at ∼241 nm.

**Fig 9.**
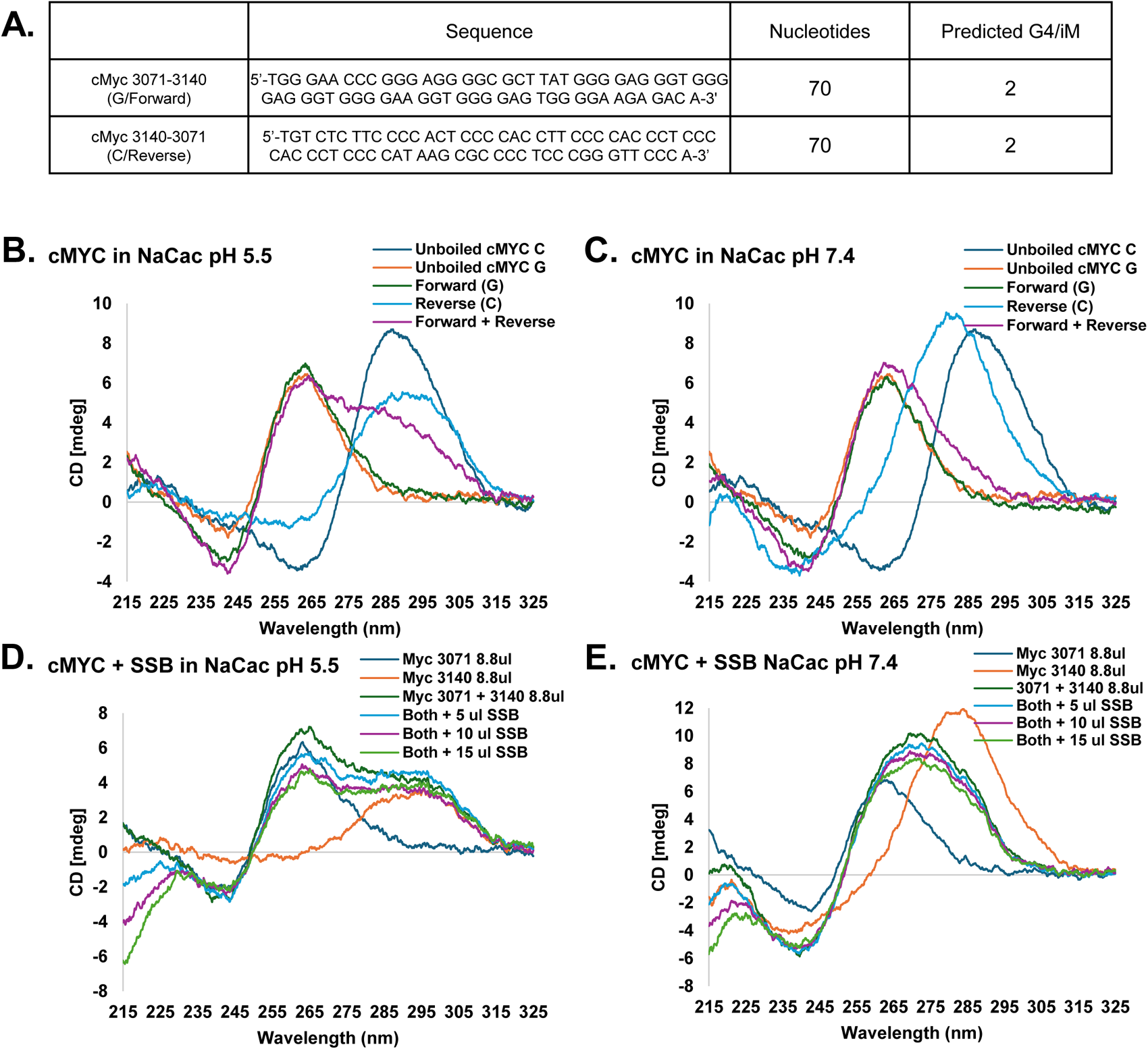
Effects of flanking sequences on G4 and iM **A.** Sequences of oligonucleotides and predictions of structure formations. **B-E.** CD Spectra of flanking sequences

**Fig. 9C** shows the same oligo combinations as **Fig. 9B**, but in the higher pH 7.4 buffer. Both unboiled cMYC samples show the same ellipticity as **Fig. 9B**. Forward (G) has the same peak wavelength as in the pH 5.5 buffer, but a slightly suppressed ellipticity. The major difference between pH 5.5 and 7.4 is the Reverse (C) oligo. While not unexpected, Reverse (C) shifted ∼10nm from ∼290nm in pH 5.5 to ∼280nm in pH 7.4. In addition to the wavelength shift, the ellipticity also had a significant increase from ∼5 mdeg to ∼9 mdeg. The combination of Forward and Reverse oligos also produced interesting results. At pH 7.4, the combination of complimentary strands eliminated the secondary i-motif peak seen in pH 5.5. The ellipticity also slightly increased but produced the conventional G-quadruplex spectra formation.

In addition to the experiments with competing complimentary sequences, CD analysis was also carried out to test the effects of SSB on dsDNA formation. As shown in **Fig. 9D**, the addition of three different concentrations of SSB affects the CD patterns when both sequences are present in the same solution. The addition of 5 ug of SSB reduced the ellipticity by ∼2 mdeg while the addition of 10 ug only reduced ellipticity by an additional ∼1 mdeg. The addition of 15 ug of SSB had almost no ellipticity change compared to the 10 ug previously added. Interestingly, the addition of SSB slightly increased the ellipticity of the i-motif peak around ∼295 nm, when compared to the Forward and Reverse complimentary strand combination.

As with the experiment in **Fig. 9 D**, SSB was added in three different concentrations to cMYC Forward and Reverse sequences in the buffer containing Na.Cac pH 7.4 (**Fig. 9 E**). Unlike the sodium cacodylate pH 5.5 buffer, there were no secondary positive peaks. Interestingly, the peaks shifted to ∼273 nm and reduced equally with each addition of SSB. This new peak formation at ∼273 nm is not consistent with either G4 or iM formation. Negative ellipticity values significantly reduced and formed a smoother curve.

## DISCUSSIONS

This study addresses the importance of multiple factors that regulate the G4- and iM-formation in C9ORF72 intron 1, and promoter-proximal regions of BCL-2, c-MYC, EGFR, HIF-1a, hRAS, and VEGF in a comparative manner. The factors include the arrangement of nucleotide sequences, DNA sequences flanking the G4/iM-forming sequences, complimentary DNA sequences, pH, buffer conditions, temperature, and SSB, a single-strand DNA binding protein which binds DNA in a sequence-independent manner. Until now, extensive studies were carried out on G4- and iM formation in c-MYC and BCL2, but with limited information available on HIF-1α, EGFR, VEGF, hRAS, and C9ORF72.

The CD patterns of tandem repeats (5’-GGGGCC-3’ and 5’-CCCCGG-3’) are different from that of DNA sequences containing polyG/polyC stretches not arranged in tandem (e.g., sequences at promoter-proximal regions of various oncogenes). The stability and CD pattern of GGGGCC and CCCCGG repeats changed as the number of the hexanucleotide repeats increased. C9ORF72 is the most defective gene in ALS and FTD^8,28,29^ in which the hexanucleotide repeat “GGGGCC/GGCCCC”, within the first intron of the gene, expands from <20 tandem copies to hundreds or thousands of tandem copies^8^. Until now, limited information was available regarding the hexanucleotide repeats. G4 formation has been studied at high temperature (85°C) in the presence of potassium cation in phosphate buffer^8^. Another study^9^ analyzed the CD spectra of “GGGGCC” repeats under varying pH (5, 5.2, 5.4, 5.6, 6), varying temperatures (15-95 °C), and the presence of different ions (lithium, potassium, magnesium, and sodium). This study also investigated the result of varying the G:C ratio of the repeat. The results had revealed an unusual, inverted CD spectrum in the presence of lithium ion. The study hypothesized that this structure might be either an unorthodox G4/iM composite, a left-handed Hoogsteen base-paired duplex or a non-canonical, “braided” DNA triplex. The structure appears to be most stable at pH 5.0-5.2, and begins to be disrupted at higher pH values, as well as in the presence of potassium cation. The melting temperature for this structure was calculated to be around 60°C, depending on the buffer. Varying the G:C concentration largely resulted in the disruption of the structure. A study using 4, 6, or 8 units of C9ORF72 C repeats shows iM formation at pH < 5, and monomolecular hairpins and monomolecular quadruplexes at pH > 8 (pH 3.95-9.24)^29^; both structures begin to denature at temperatures above 60°C

BCL-2 transcription^14^ is regulated by P1 and P2 promoters of which the P1 promoter containing G4- and iM-forming sequences is responsible for the majority of BCL-2 transcription^25^. At P1, G4 persists at a temperature range of 20-82 °C in 10 mM Tris–HCl/150 mM KCl (pH 7.2), which is destabilized by Li+ cation^25^. Another study^39^ investigated equimolar mixtures of the G- and C-rich strands of the BCL-2 promotor, exploring pH dependence, temperature effects, and the influence of potassium and cesium ions. Results showed that at low potassium ion concentrations, BCL-2 predominantly adopts a duplex conformation, with approximately 10% of sequences forming iM; however, at high concentrations of potassium ions there was coexistence of the duplex, G4s, and iMs. Another study has addressed iM formation across varying pH and temperature conditions at the P1 promoter^40^. A study using 50 mM Tris buffer (pH 6.5 or 7.5) and 100 mM NaCl has addressed the interaction of the iM-forming sequence with hnRNP LL protein^41^. Lastly, a study has shown the prevalence of iMs over the hairpin structure at pH values less than 7 at 10°C sodium phosphate buffer^42^. c-MYC gene, which regulates transcription of ∼15% of human genes, has been implicated in ∼20% of all cancers^33^. CD analysis of G4 at the NHE III1 region of c-MYC has been studied extensively. A CD analysis of c-MYC G4 formation using buffers including 10 mM TBA acetate (pH 5.0), 10 mM TBA cacodylate (pH 6.0), and 10 mM TBA phosphate (pH 7.0) across a wide range of temperatures (20-100 °C), pHs (5.0, 6.0, or 7.0), and concentrations of KCl (0-50 mM) has shown that whereas high concentrations of KCl promote G4, low KCl concentrations do not do so^43^. Various ligands have been developed that were shown to stabilize formation of c-MYC G4 across a wide range of temperatures (0-100°C) in 10 mM Tris-HCl buffer (pH 7.2)^44^. The protein Nucleolin also has also been shown to stabilize such G4 formation in 50 mM Tris-HCl (pH 7.4)^45^. Study of iM at c-MYC has been studied less extensively. Although iM formation is lost at pH 6.0 or higher pH, molecular crowding by high molecular weight polyethylene glycol (PEG_8000_) can stabilize iMs at pH values as high as 6.7 in BPES buffer^2.4^. It has been demonstrated that addition of a molecular crowding agent Namely Pyrene can stabilize c-MYC iM in 10 mM Tris buffer ( pH 5.5-7.5) at temperatures 10°C higher than that without it in a range of 20-80°C. HIF-1α plays a key role in the expression of many genes involved in metabolic pathways including angiogenesis, inflammatory response to hypoxia, anaerobic metabolism and tumor proliferation^17^. HIF1- α is expressed in response to hypoxic states, growth factor expression, and carcinogen expression with the goal of promoting cellular survival via angiogenesis and adaptation to anaerobic metabolism whereas HIF1-Beta is constitutively active^47^. However, information on CD patterns of G4s and iMs at HIF-1α promoter is not elaborate. G4 is formed in the promotor region of the HIF-1α subunit^48^, and it has been suggested that stable iM can form in the region^49^. The polyG/polyC-rich sequences at the promoter of EGFR gene can form two G4s (one parallel and one hybrid) in 10 mM Tris buffer (pH 7.5) across a range of temperatures (20-100 °C) and KCl concentrations (0-200 mM) and specific mutations abolished the structures^50^. A CD analysis has been carried out on the polyC-rich DNA in EGFR promoter across a range of pH values (5.5 – 8.6), in the presence of varying concentration of KCl (0–250 mM), and different crowding agents (20%-40% PEG_300_) on iM in the same region^51^.. Results indicate that iM was formed in the promotor region of EGFR, and that at neutral pH, iM and hairpin structures coexist; 10 mM Tris pH 7.0 buffer was used for this experiment. The two polyC-rich regions of hRAS (hRAS1 and hRAS2) have been analyzed by CD spectroscopy for iM generation under a range of pH values (4.5 – 8.0), temperatures (20-80°C), and the polyC-binding protein hnRNP A1 in 50 mM Tris-acetate, 50 mM KCl, 40% PEG-300 buffer (pH 4.5 – 8.0) or 50 mM sodium cacodylate, 50 mM KCl (pH 5.0)^20^. hnRNP A1 unfolded the iMs. In another study CpG methylation was found to stabilize iMs and marginally elevate the melting temperature^52^. CD analysis has shown an increase in G4 peak with increasing concentration of KCl when experiments were carried in buffers containing a range of KCl concentrations (0 - 100 mM) in 20 mM Tris–HCl buffer (pH 7.6) at 25°C^53^. Chelerythrine, an anticancer plant alkaloid, has been found to stabilize G4 at higher temperature in buffers containing10 mM potassium phosphate buffer with 100 mM KCl (pH 7.0)^54^. Doxorubicin, another anticancer agent, also stabilizes G4 structure at high temperature (95°C) in K-phosphate buffer with 70 mM KCl, pH of 7.0^55^. Formation of iM occurs at VEGF promoter region and it decreased as the pH increased from 4.4 to 8.0^56^. A study that investigated both G4 and iM formation within the VEGF gene in 10 mM TBA phosphate buffer under different pH values (3.31 – 9.80), KCl concentrations (0 – 50 mM), and temperatures (25 - 95 °C) has shown that at pH 5.0 or 7.0, room temperature, and low concentrations of potassium ions, a majority of DNA exists in the duplex state with about 10% forming iM^39^. However, at higher concentrations of potassium ions, G4, iM, and duplex structures all coexist.

Most of the above-mentioned CD analyses were carried out with individual oligos in isolation and under specific conditions. The present study was carried out with all oligos under the same conditions, revealing variations among the G4s and iMs. Thus, the current work can be used as a ready reference for G4 and iM formation by all these oligos. Oligos containing small number of hexanucleotide repeats of C9ORF72 intron 1 display iMs and G4s as expected; however, as the repeat number increases, CD spectra do not follow the expected patterns with changes in pH. We predict that G:C base-pairing becomes more prevalent to cause these variations. CD spectra after boiling and cooling the oligos displayed significant differences from the unboiled counterparts in some, but not all oligos. We ascribe these differences to the DNA sequence in the oligos. One of the surprising and interesting findings is that boiling and cooling of a population of two complementary oligos does not convert the entire population into dsDNA. The presence of flanking sequences affect G4 and iM patterns of c-MYC promoter-proximal DNA sequences when both oligos are mixed in solution. Finally, *E. coli* SSB reduces the population of both G4s and iMs in almost all oligos tested.

## MATERIALS AND METHODS

### Buffers

Buffers used are Na.Cac pH 5.5 (10 mM sodium cacodylate, 100 mM KCl), Tris.acetate pH 6.0 (Tris.acetate 20 mM, 100 mM KCl), MES pH 6.5 (MES 50 mM, 100 mM, MgCl_2_ 2 mM, EDTA 1 mM, DTT 1 mM, Tween20 0.1%, glycerol 10%, Na.Cac pH 7.4, 100 mM KCl), Tris.KCl pH 7.4 (Tris 50 mM, KCl 100 mM).

### DNA Oligodeoxynucleotides (oligos)

All oligodeoxynucleotides were obtained commercially from Millipore or IDT. Oligos were HPLC purified, shipped dry and reconstituted with TE (10 mM Tris pH 7.4, 1 mM EDTA pH 8.0). See Supplementary Table 1 for sequences of the oligos.

### Protein

*E. coli* single-strand DNA binding protein (SSB) and hnRNP K were obtained commercially from MCLAB and LSBio, respectively.

### Circular dichroism (CD) measurements

All spectra were measured using a Jasco J-1500 circular dichroism spectrophotometer with either the standard adjustable cell holder or the single-position Peltier thermostatted cell holder. Non-temperature-controlled samples were measured in a 1mm path length cell and temperature-controlled samples were measured in a 10mm path length cell. Spectra were obtained using the following parameters: 0.1 nm data pitch, 50 nm/min scanning speed, 200 mdeg/0.1 dOD, 2 sec D.I.T and 1 nm bandwidth.

### Polyacrylamide gel electrophoresis (PAGE) and urea-polyacrylamide gel electrophoresis (urea-PAGE)

All oligos were fractionated by 10% PAGE or 15% urea-PAGE, stained with red dye, and analyzed by iBright 1500 imager (Invitrogen).

## ACKNOWLEDGEMENTS

BKM was supported by VCOM REAP grants (#1032453 and #1302559).

## Supplementary

**Table S1.**
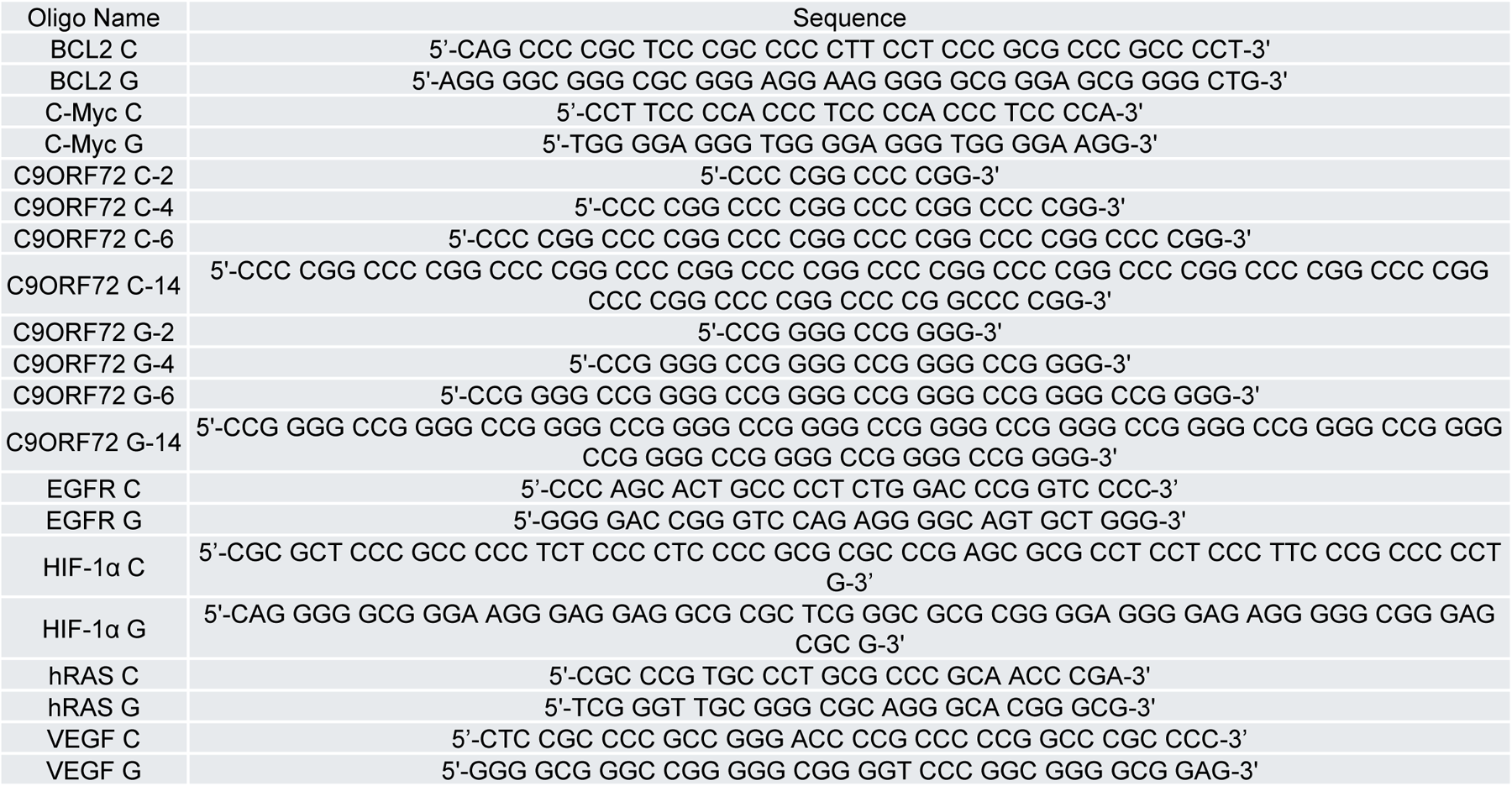
Sequences of Oligodeoxynucleotides.

**Table S2.**
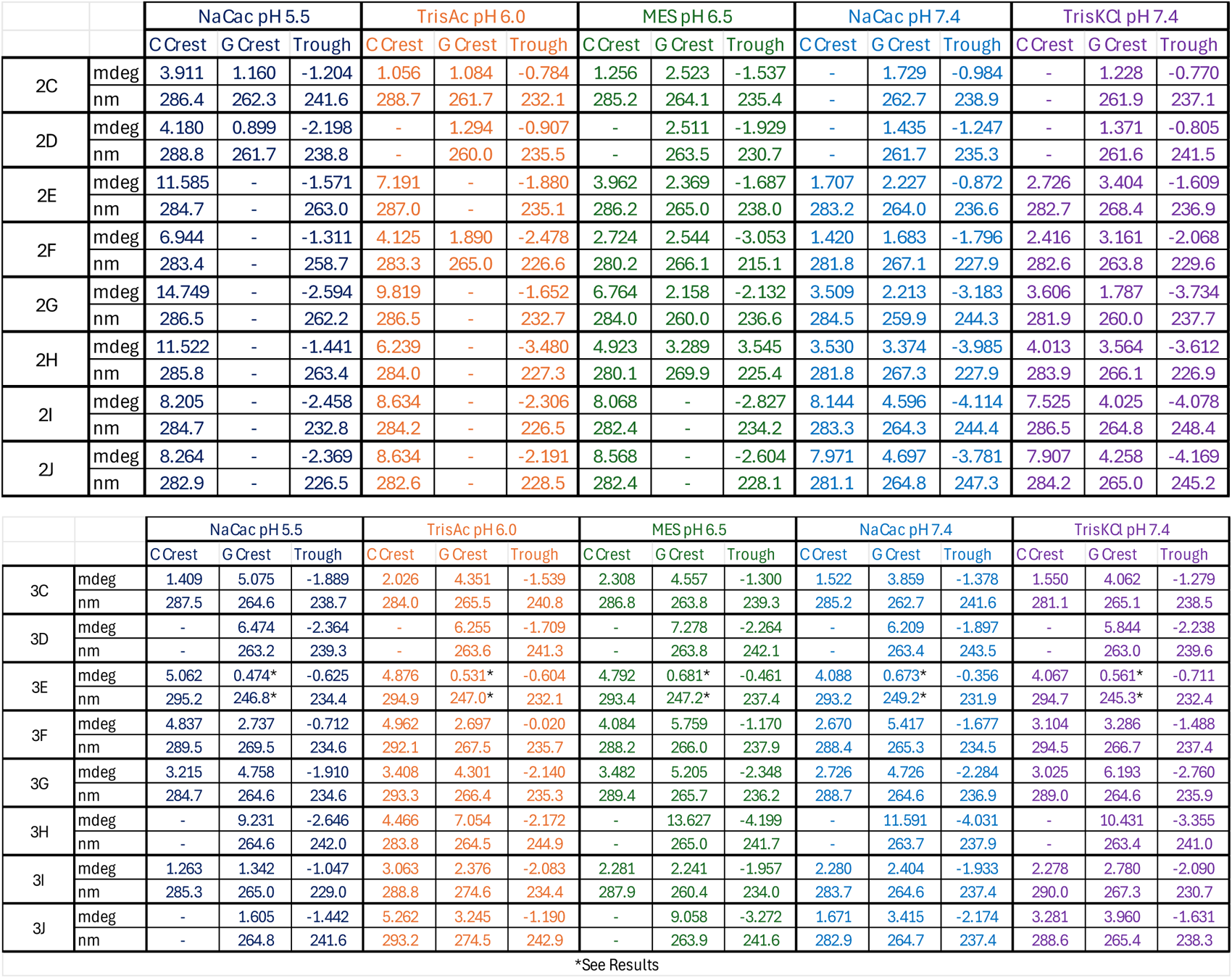
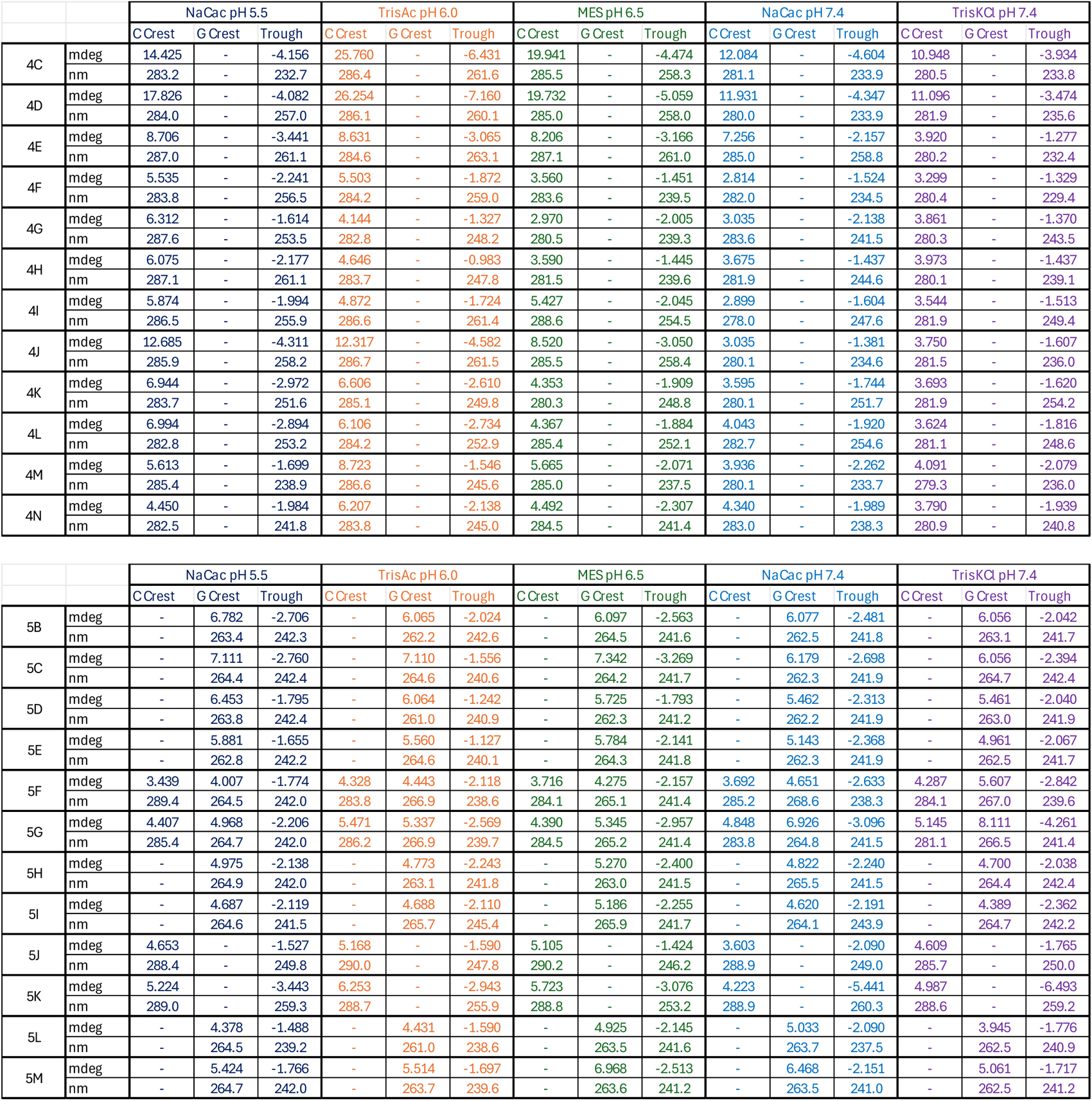
Peak Values of iM and G4.

## REFERENCES

1. Poggi, L. & Richard, G.-F. Alternative DNA Structures In Vivo: Molecular Evidence and Remaining Questions. Microbiol. Mol. Biol. Rev. MMBR 85, (2021).

2. Kaushik, M., Kaushik, S., Bansal, A., Saxena, S. & Kukreti, S. Structural diversity and specific recognition of four stranded G-quadruplex DNA. Curr. Mol. Med. 11, 744–769 (2011).

3. Ortiz de Luzuriaga, I., Lopez, X. & Gil, A. Learning to Model G-Quadruplexes: Current Methods and Perspectives. Annu. Rev. Biophys. 50, 209–243 (2021).

4. Alba, J. J., Sadurní, A. & Gargallo, R. Nucleic Acid i-Motif Structures in Analytical Chemistry. Crit. Rev. Anal. Chem. 46, 443–454 (2016).

5. Chambers, V. S. et al. High-throughput sequencing of DNA G-quadruplex structures in the human genome. Nat. Biotechnol. 33, 877–881 (2015).

6. Peña Martinez, C. D., et al. Human genomic DNA is widely interspersed with i-motif structures. bioRxiv (2022).

7. Balendra, R. & Isaacs, A. M. C9orf72-mediated ALS and FTD: multiple pathways to disease. Nat. Rev. Neurol. 14, 544–558 (2018).

8. Fratta, P. et al. C9orf72 hexanucleotide repeat associated with amyotrophic lateral sclerosis and frontotemporal dementia forms RNA G-quadruplexes. Sci. Rep. 2, 1016 (2012).

9. Lat, P. K. & Sen, D. (C2G4) n repeat expansion sequences from the C9orf72 gene form an unusual DNA higher-order structure in the pH range of 5-6. PLoS One 13, e0198418 (2018).

10. Valton, A.-L. & Prioleau, M.-N. G-Quadruplexes in DNA Replication: A Problem or a Necessity? Trends Genet. TIG 32, 697–706 (2016).

11. Bay, D. H. et al. Identification of G-quadruplex structures that possess transcriptional regulating functions in the Dele and Cdc6 CpG islands. BMC Mol. Biol. 18, 17 (2017).

12. Chen, L., Dickerhoff, J., Sakai, S. & Yang, D. DNA G-Quadruplex in Human Telomeres and Oncogene Promoters: Structures, Functions, and Small Molecule Targeting. Acc. Chem. Res. 55, 2628–2646 (2022).

13. Dvoráková, Z. et al. i-Motif of cytosine-rich human telomere DNA fragments containing natural base lesions. Nucleic Acids Res. 46, 1624–1634 (2018).

14. Frenzel, A., Grespi, F., Chmelewskij, W. & Villunger, A. Bcl2 family proteins in carcinogenesis and the treatment of cancer. Apoptosis 14, 584–596 (2009).

15. Wang, W. et al. Human MYC G-quadruplex: From discovery to a cancer therapeutic target. Biochim. Biophys. Acta Rev. Cancer 1874, 188410 (2020).

16. Dai, J., Hatzakis, E., Hurley, L. H. & Yang, D. I-motif structures formed in the human c-MYC promoter are highly dynamic--insights into sequence redundancy and I-motif stability. PloS One 5, e11647 (2010).

17. Shi, Y.-H. & Fang, W.-G. Hypoxia-inducible factor-1 in tumour angiogenesis. World J. Gastroenterol. 10, 1082 (2004).

18. Normanno, N. et al. Epidermal growth factor receptor (EGFR) signaling in cancer. Gene 366, 2–16 (2006).

19. Shibuya, M. Vascular endothelial growth factor (VEGF) and its receptor (VEGFR) signaling in angiogenesis: a crucial target for anti-and pro-angiogenic therapies. Genes Cancer 2, 1097–1105 (2011).

20. Miglietta, G., Cogoi, S., Pedersen, E. B. & Xodo, L. E. GC-elements controlling HRAS transcription form i-motif structures unfolded by heterogeneous ribonucleoprotein particle A1. Sci. Rep. 5, 18097 (2015).

21. Santra, T. et al. An integrated global analysis of compartmentalized HRAS signaling. Cell Rep. 26, 3100–3115 (2019).

22. Fratta, P. et al. C9orf72 hexanucleotide repeat associated with amyotrophic lateral sclerosis and frontotemporal dementia forms RNA G-quadruplexes. Sci. Rep. 2, 1016 (2012).

23. Kovanda, A., Zalar, M., Šket, P., Plavec, J. & Rogelj, B. Anti-sense DNA d(GGCCCC)n expansions in C9ORF72 form i-motifs and protonated hairpins. Sci. Rep. 5, 17944 (2015).

24. Murat, P., Singh, Y. & Defrancq, E. Methods for investigating G-quadruplex DNA/ligand interactions. Chem. Soc. Rev. 40, 5293–5307 (2011).

25. Sun, H. et al. A newly identified G-quadruplex as a potential target regulating Bcl-2 expression. Biochim. Biophys. Acta BBA-Gen. Subj. 1840, 3052–3057 (2014).

26. Shetu, S. A. & Bandyopadhyay, D. Small-molecule RAS inhibitors as anticancer agents: discovery, development, and mechanistic studies. Int. J. Mol. Sci. 23, 3706 (2022).

27. Aoki, Y. et al. Germline mutations in HRAS proto-oncogene cause Costello syndrome. Nat. Genet. 37, 1038–1040 (2005).

28. Smeyers, J., Banchi, E. & Latouche, M. C9ORF72: what it is, what it does, and why it matters. Front Cell Neurosci 15: 661447. (2021).

29. Zamiri, B. et al. Stress-induced acidification may contribute to formation of unusual structures in C9orf72-repeats. Biochim. Biophys. Acta BBA-Gen. Subj. 1862, 1482–1491 (2018).

30. Kendrick, S. & Hurley, L. H. The role of G-quadruplex/i-motif secondary structures as cis-acting regulatory elements. Pure Appl. Chem. Chim. Pure Appl. 82, 1609–1621 (2010).

31. Yu, H. et al. iM-Seeker: a webserver for DNA i-motifs prediction and scoring via automated machine learning. Nucleic Acids Res. gkae315 (2024) doi:10.1093/nar/gkae315.

32. Del Villar-Guerra, R., Trent, J. O. & Chaires, J. B. G-Quadruplex Secondary Structure Obtained from Circular Dichroism Spectroscopy. Angew. Chem. Int. Ed Engl. 57, 7171–7175 (2018).

33. Dang, C. V. et al. The c-Myc target gene network. in vol. 16 253–264 (Elsevier, 2006).

34. Antony, E. & Lohman, T. M. Dynamics of E. coli single stranded DNA binding (SSB) protein-DNA complexes. Semin. Cell Dev. Biol. 86, 102–111 (2019).

35. Dueva, R. & Iliakis, G. Replication protein A: a multifunctional protein with roles in DNA replication, repair and beyond. NAR Cancer 2, zcaa022 (2020).

36. Chowdhury, S., Chowdhury, A. B., Kumar, M. & Chakraborty, S. Revisiting regulatory roles of replication protein A in plant DNA metabolism. Planta 253, 130 (2021).

37. Sun, Z.-Y., Wang, X.-N., Cheng, S.-Q., Su, X.-X. & Ou, T.-M. Developing Novel G-Quadruplex Ligands: from Interaction with Nucleic Acids to Interfering with Nucleic Acid^−^Protein Interaction. Mol. Basel Switz. 24, (2019).

38. Wu, Y. & Brosh, R. M. J. G-quadruplex nucleic acids and human disease. FEBS J. 277, 3470–3488 (2010).

39. Liu, L. et al. Distribution of Conformational States Adopted by DNA from the Promoter Regions of the VEGF and Bcl-2 Oncogenes. J. Phys. Chem. B 126, 6654–6670 (2022).

40. Kendrick, S., Akiyama, Y., Hecht, S. M. & Hurley, L. H. The i-motif in the bcl-2 P1 promoter forms an unexpectedly stable structure with a unique 8: 5: 7 loop folding pattern. J. Am. Chem. Soc. 131, 17667–17676 (2009).

41. Basab, R., Poulami, T. & Hyun-Jin, K. Interaction of Individual Structural Domains of hnRNP LL with the BCL2 Promoter i-Motif DNA. (2016).

42. Amato, J. et al. Conformational plasticity of DNA secondary structures: Probing the conversion between i-motif and hairpin species by circular dichroism and ultraviolet resonance Raman spectroscopies. Phys. Chem. Chem. Phys. 24, 7028–7044 (2022).

43. Liu, L., Ma, C., Wells, J. W. & Chalikian, T. V. Conformational Preferences of DNA Strands from the Promoter Region of the c-MYC Oncogene. J. Phys. Chem. B 124, 751–762 (2020).

44. Głuszyńska, A., Juskowiak, B., Kuta-Siejkowska, M., Hoffmann, M. & Haider, S. Carbazole ligands as c-myc G-quadruplex binders. Int. J. Biol. Macromol. 114, 479–490 (2018).

45. González, V. & Hurley, L. H. The C-terminus of nucleolin promotes the formation of the c-MYC G-quadruplex and inhibits c-MYC promoter activity. Biochemistry 49, 9706–9714 (2010).

46. Cui, J., Waltman, P., Le, V. H. & Lewis, E. A. The effect of molecular crowding on the stability of human c-MYC promoter sequence I-motif at neutral pH. Molecules 18, 12751– 12767 (2013).

47. Ziello, J. E., Jovin, I. S. & Huang, Y. Hypoxia-Inducible Factor (HIF)-1 regulatory pathway and its potential for therapeutic intervention in malignancy and ischemia. Yale J. Biol. Med. 80, 51 (2007).

48. De Armond, R., Wood, S., Sun, D., Hurley, L. H. & Ebbinghaus, S. W. Evidence for the presence of a guanine quadruplex forming region within a polypurine tract of the hypoxia inducible factor 1α promoter. Biochemistry 44, 16341–16350 (2005).

49. Brazier, J. A., Shah, A. & Brown, G. D. I-motif formation in gene promoters: unusually stable formation in sequences complementary to known G-quadruplexes. Chem. Commun. 48, 10739–10741 (2012).

50. Greco, M. L. et al. Coexistence of two main folded G-quadruplexes within a single G-rich domain in the EGFR promoter. Nucleic Acids Res. 45, 10132–10142 (2017).

51. Cristofari, C., Rigo, R., Greco, M. L., Ghezzo, M. & Sissi, C. pH-driven conformational switch between non-canonical DNA structures in a C-rich domain of EGFR promoter. Sci. Rep. 9, 1210 (2019).

52. Oshikawa, D., Inaba, S., Kitagawa, Y., Tsukakoshi, K. & Ikebukuro, K. CpG methylation altered the stability and structure of the i-motifs located in the CpG islands. Int. J. Mol. Sci. 23, 6467 (2022).

53. Sun, D., Guo, K., Rusche, J. J. & Hurley, L. H. Facilitation of a structural transition in the polypurine/polypyrimidine tract within the proximal promoter region of the human VEGF gene by the presence of potassium and G-quadruplex-interactive agents. Nucleic Acids Res. 33, 6070–6080 (2005).

54. Jana, J. et al. Chelerythrine down regulates expression of VEGFA, BCL2 and KRAS by arresting G-Quadruplex structures at their promoter regions. Sci. Rep. 7, 40706 (2017).

55. Bilgen, E. & Çetinkol, Ö. P. Doxorubicin exhibits strong and selective association with VEGF Pu22 G-quadruplex. Biochim. Biophys. Acta BBA-Gen. Subj. 1864, 129720 (2020).

56. Guo, K., Gokhale, V., Hurley, L. H. & Sun, D. Intramolecularly folded G-quadruplex and i-motif structures in the proximal promoter of the vascular endothelial growth factor gene. Nucleic Acids Res. 36, 4598–4608 (2008).

